# Stochastic modelling of a three-dimensional glycogen granule synthesis and impact of the branching enzyme

**DOI:** 10.1101/2022.10.31.514469

**Authors:** Yvan Rousset, Oliver Ebenhöh, Adélaïde Raguin

**Affiliations:** Institute for Quantitative and Theoretical Biology, Heinrich-Heine University, Düsseldorf, Germany; Cluster of Excellence on Plant Sciences (CEPLAS), Heinrich-Heine University, Düsseldorf, Germany; Institute for Computational Cell Biology, Heinrich-Heine University, Düsseldorf, Germany

## Abstract

In humans, glycogen storage diseases result from metabolic inborn errors, and can lead to severe phenotypes and lethal conditions. Besides these rare diseases, glycogen is also associated to widely spread societal burdens such as diabetes. Glycogen is a branched glucose polymer synthesised and degraded by a complex set of enzymes. Over the past 50 years, the structure of glycogen has been intensively investigated. Yet, the interplay between glycogen structure and the related enzymes is still to be characterised. In this article, we develop a stochastic coarse-grained and spatially resolved model of branched polymer biosynthesis following a Gillespie algorithm. Our study largely focusses on the role of the branching enzyme, and first investigates the properties of the model with generic parameters, before comparing it to *in vivo* experimental data in mice. It arises that the ratio of glycogen synthase over branching enzyme activities drastically impacts the structure of the granule. We deeply investigate the mechanism of branching and parametrise it using distinct lengths. Not only do we consider various possible sets of values for these lengths, but also distinct rules to apply them. We show how combining them finely tunes glycogen macromolecular structure. Comparing the model with experimental data confirms that we can accurately reproduce glycogen chain length distributions in wild type mice. Additional granule properties obtained for this fit are also in good agreement with typically reported values in the experimental literature. Nonetheless, we find that the mechanism of branching must be more flexible than usually reported. Overall, we demonstrate that the chain length distribution is an imprint of the branching activity and mechanism. Our generic model and methods can be applied to any glycogen data set, and could in particular contribute to characterise the mechanisms responsible for glycogen storage disorders.

**Author summary:** Glycogen is a granule-like macromolecule made of 10,000 to 50,000 glucose units arranged in linear and branched chains. It serves as energy storage in many species, including humans. Depending on physiological conditions (hormone concentrations, glucose level, etc.) glycogen granules are either synthesised or degraded. Certain metabolic disorders are associated to abnormal glycogen structures, and structural properties of glycogen might impact the dynamics of glucose release and storage. To capture the complex interplay between this dynamics and glycogen structural properties, we propose a computational model relying on the random nature of biochemical reactions. The granule is represented in three dimensions and resolved at the glucose scale. Granules are produced under the action of a complex set of enzymes, and we mostly focus on those responsible for the formation of new branches. Specifically, we study the impact of their molecular action on the granule structure. With this model, we are able to reproduce structural properties observed under certain *in-vivo* conditions. Our biophysical and computational approach complements experimental studies and may contribute to characterise processes responsible for glycogen related disorders.

## 1 Introduction

Management of energy resources is crucial for an organism to survive, since nutrient availability may vary considerably with time. Moreover, organisms face numerous other stresses that may temporarily increase energy demand. To react to such changes in energy supply and demand, it is apparent that some internal stores are necessary. These stores are filled when nutrients are abundant and depleted when demand exceeds available supply. As direct substrate for catabolic pathways, glucose plays a central role in energy metabolism in most organisms [1]. While plants have found their way to store glucose as starch, animals, fungi, and most bacteria store glucose as glycogen. Both starch and glycogen are branched polymers consisting of glucose monomers, linked into linear chains by *α*-1,4 bonds, and branching points by *α*-1,6 bonds. However, these two macromolecules exhibit rather different physico-chemical properties. In contrast to glycogen, starch is insoluble in water under physiological conditions, and contains high density crystalline regions. These different properties are reflected by distinct branching patterns and chain length distributions. Functionally, starch and glycogen can be compared to capacitors in electric circuits. The latter are able to store and release electrons depending on current and voltage. Thus, they can be used to stabilise a fluctuating electric signal. Analogously, glycogen and starch can be seen as two different capacitors that both contribute to glucose homeostasis by managing energy resources through time.

While the fine structure of starch has been widely investigated over the past 50 years, less is known on that of glycogen [2]. The characterisation of the detailed structure of glycogen, as well as the interplay between its structural properties and the dynamics of glycogenesis (synthesis) and glycogenolysis (degradation) is unclear. Yet, both glycogen structure and dynamics are of utmost interest for understanding glycogen metabolism and the impact of related genetic variances. For human health, this is particularly important considering the increasing prevalence of glycogen storage diseases (GSDs), as well as diabetes, and other glycogen related disorders.

So far, different structures of glycogen have been proposed [3–6], but a structural model that emerges from the detailed underlying enzymatic mechanisms of synthesis and degradation is still lacking. Understanding and precisely describing the biochemistry of glycogen is challenging. With a molecular weight of 10^6^ to 10^7^ g·mol^−1^ [7–9], glycogen is a large molecule, even compared to the enzymes involved in its dynamics [10–12]. Thus, enzymes synthesising, degrading, or otherwise altering glycogen, can only access certain branches near its surface, while many glucose residues near the centre of the molecule remain ‘hidden’. At the macroscopic scale, the structure of glycogen is well known. Drochmans [13] observed two populations of glycogen granules in rat liver. The so-called *α* granules are aggregates of the smaller *β* granules [14, 15]. The latter have a radius in the range from 10 to 20 nm, while the radius of *α* granules ranges between 40 and 300 nm [13]. Here, we focus on *β* granules, whose synthesis and degradation both involve a relatively small number of enzymes (see Fig 1). For a comprehensive review, see [16].

**Fig 1.**
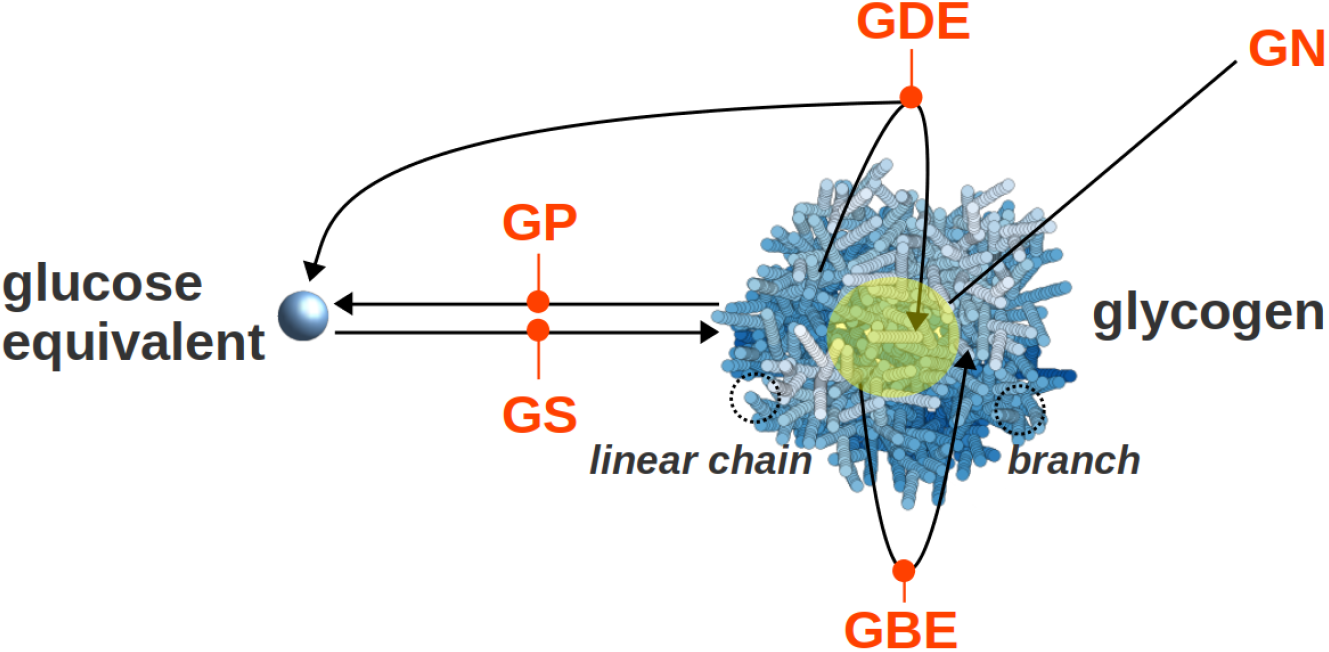
Main enzyme reactions involved in the synthesis and breakdown of glycogen. *In vivo*, the GS and GBE enzymes synthesise glycogen, while the GP and GDE degrade it. Besides, GN is the initial precursor of the granule and stands in its core. Enzymes are noted in orange, glucose residues are in blue, and GN is highlighted with a yellow sphere.

To initiate a new glycogen molecule, a chain of 5 glucose units is synthesised and bound to a glycogenin (GN) protein, which forms the core of the final granule [17–20]. Once initiation is completed, the granule is expanded by the two enzymes Glycogen Synthase (GS) and Glycogen Branching Enzyme (GBE). GS is an elongating enzyme that adds one glucose residue to the non-reducing end of an *α*-1,4 linear chain, thereby forming a new *α*-1,4 glucosydic bond. GBE cleaves a small part of a linear chain and creates a new branch by forming an *α*-1,6 glucosydic bond. We call “daughter” the newly formed chain and “mother” the one it is branching from.

Besides synthesis, granules are subject to degradation, that is performed by two other enzymes. Glycogen Phosphorylase (GP) and Glycogen Debranching Enzyme (GDE) respectively shorten and debranch glycogen branches.

Depending on the relative enzymatic activities of these four enzymes (GS, GBE, GP, and GDE) the overall size of the glycogen granule can either increase or decrease. In this article, we choose to focus on glycogen synthesis, and more specifically the role of the branching enzyme GBE.

Experimental observations [21] of glycogen show an average chain length (CL) of 13 glucose units. The chain length distribution is not symmetric [22, 23]. A typical peak is observed at low degree of polymerisation (DP), around DP 8, and almost no chains are detected above DP 40. The degree of branching is defined in two ways in the literature. It is most commonly defined as the ratio of *α*–1,6 to *α*–1,4 linkages, but sometimes also as the average number of *α*–1,6 bonds per chain [6]. We will apply the first definition throughout the paper. This ratio is in the range 0.02 – 0.05 in amylopectin [24–26] from starch and 0.06 – 0.10 in glycogen [9, 27]. In 1956, Peat et al. [28] introduced the concept of A and B chains. An A chain does not carry any branch, while a B chain does. The A:B ratio is an important characteristic of glycogen granules and an indicator of the branching pattern. Early studies reported an A:B ratio of 1 in glycogen [21,29], while it is usually greater than 1 in amylose, and ranging from 1.5 to 2.6 in amylopectine [29, 30].

As early as in the 1940s various hypotheses have been formulated aiming at explaining macroscopic features of glycogen granules. One of them has become known as the Whelan model, which assumes that every chain carries on average two branches. Based on the Whelan model, the nowadays widespread idea emerged that glycogen can be described as a fractal molecule [6, 31, 32]. A fractal glycogen structure is indeed attractive because it can reproduce various structural properties of glycogen. Moreover, it provides a mechanism explaining why glycogen granules seem to have a maximum size. With this model, glycogen becomes too dense at the surface due to an exponential increase of the number of non-reducing ends with the distance from the centre. Thus, steric hindrance prevents synthesis to continue. However, Manners [2] stressed in 1990 that there is no clear evidence that supports a regular branching and therefore a fractal pattern. More recently, further arguments and results against a fractal structure have been raised [33, 34]. Besides, independently from any fractal structure considerations, Zhang et al. [35] proposed a mathematical model based on a Monte Carlo approach to numerically simulate glycogen biosynthesis, aiming to support the common idea that steric hindrance limits granule growth. While their model indeed shows limited growth, this results from a specific choice of parameter values, that is not supported by experimental evidences.

In this article, we propose a mechanistic model for glycogen synthesis with a major focus on the impact of the branching enzyme on the granule structure. We aim to explain macroscopic and experimentally observable quantities from the underlying enzymatic mechanisms. We first detail the geometrical and structural features, and enzymatic reactions, taken into consideration in our model. Then, we analyse distinct properties of the model with a specific focus on the effect of the branching enzyme. Finally, we compare the model to experimental data and discuss the parameter values resulting in a best fit, in relation to typical values reported in the experimental literature. In addition, several complementary results justifying our modelling choices are reported in the extensive Supplementary Material.

## 2 Results

### 2.1 The model

#### Glycogen structure and geometry

We represent glycogen granules as simplified three-dimensional structures, in which every glucose monomer is characterised by its position in space (see Fig 2). To describe the branched tree-like structure, we generalise the simple representation of linear self-avoiding polymers. Using X-ray experiments, Goldsmith et al [31] characterised in detail how glucose molecules are arranged into helical *α*-1,4 linear chains. The cross-section of the helix has been calculated to be 1.3 nm^2^ with 6 to 7 residues per turn of length 1.4 nm. The radius of the circular cross-section is thus *ρ* = 0.65 nm, and each glucose residue contributes to the chain length by *l* = 0.24 nm. Inspired by these findings, we propose that monomers are described as overlapping spheres with radius *ρ* = 0.65 nm, equal to that of the helix. The validity of this assumption and its impact on our results are presented in detail in the Supplementary Material (see Fig 11). Besides, the center of consecutive monomers are distant by *l* = 0.24 nm, which corresponds to the contribution of one glucose unit to the chain length, but also involves that the coarse-grained monomer spheres overlap. However, to realistically reflect self-avoidance, two different chains cannot. With these spatial considerations, we ensure that the contribution to the volume by one glucose unit in the model is similar to that of the real helical chain. This way, we provide a description which is simple enough to be easily represented in a computer model, but still realistic enough to reflect the spatial properties of linear chains arranged into helices.

**Fig 2.**
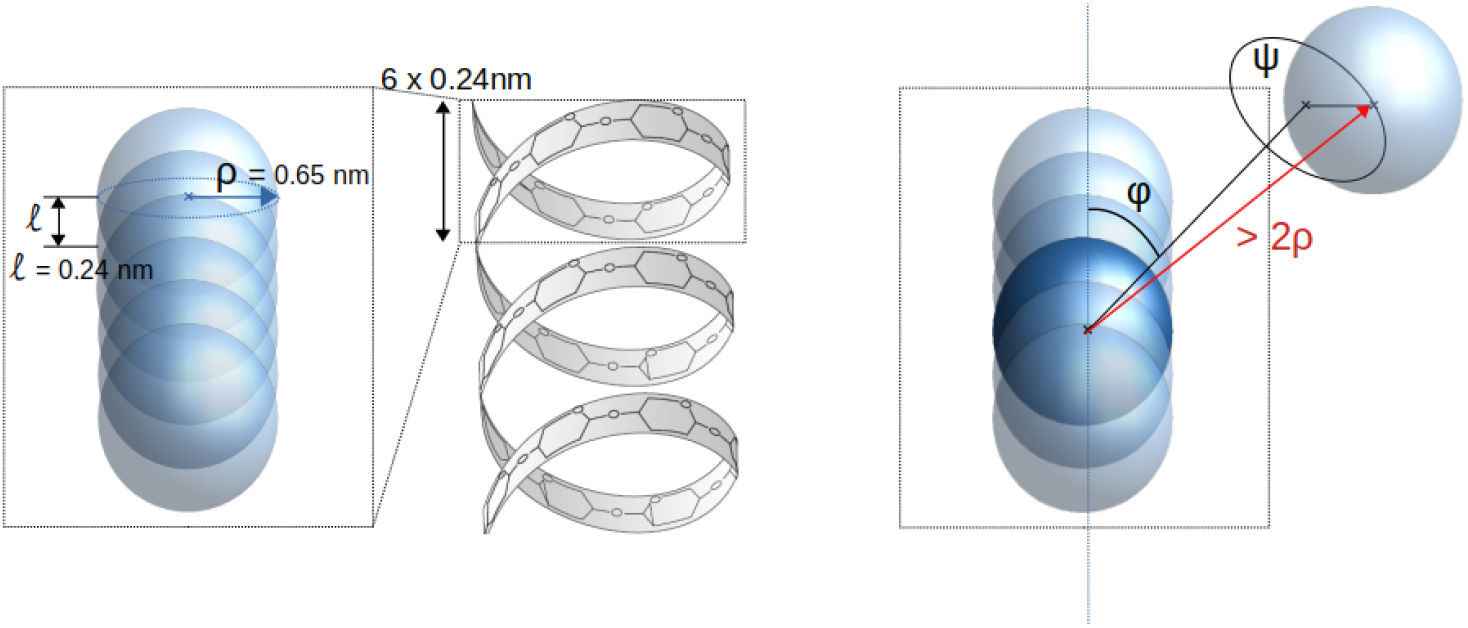
Geometrical description of glucose chains. Left: Coarse-grained linear chain. Assuming helical chains, glucose units are described as interpenetrated spheres with radius *ρ* = 0.65 nm. Two consecutive glucoses are distant by *l* = 0.24 nm, which is the radial contribution to the chain length of one glucose in a helical structure. **Right: Description of a branching point.** We generate the direction of the new branch by randomly picking two angles *φ* and *ψ*. The first monomer of the new branch will be located at a distance greater than 2*ρ* to insure no overlapping between the mother and the daughter branches.

Describing branches formed by *α*-1,6 linkages requires additional geometrical considerations. As illustrated in Fig 2 (right panel), a branch point is defined by the monomer on the mother chain to which it is attached, and two angles defining the direction of the daughter chain. Besides, the anchoring monomer on the mother chain and the first one of the daughter chain are distant by 2*ρ*, ensuring that they do not overlap.

Glycogen synthesis is initiated by GN that is located in the centre of the molecule [36]. It contributes to the total volume of glycogen, and thus we also consider self-exclusion between GN and any glucose residue of the granule. For the sake of simplicity, we assume that it is a sphere of density 1.3 g · cm^−3^ [37], which is the typical density of a protein. Accounting for its two sub-units of 38 kDa each [11], we approximate its radius by *ρ*_GN_ ≈ 2.85 nm. Together with GS, which is responsible for elongation, GN may initiate more than a single primary chain, possibly either 2 or 4 [20]. We model this initial core structure with two primary chains pointing out of the GN sphere in opposite directions.

##### 2.1.1 Enzymatic reactions

Two enzymes (GS and GBE) are directly involved in glycogen synthesis. Their role is illustrated in Fig 3.

**Fig 3.**
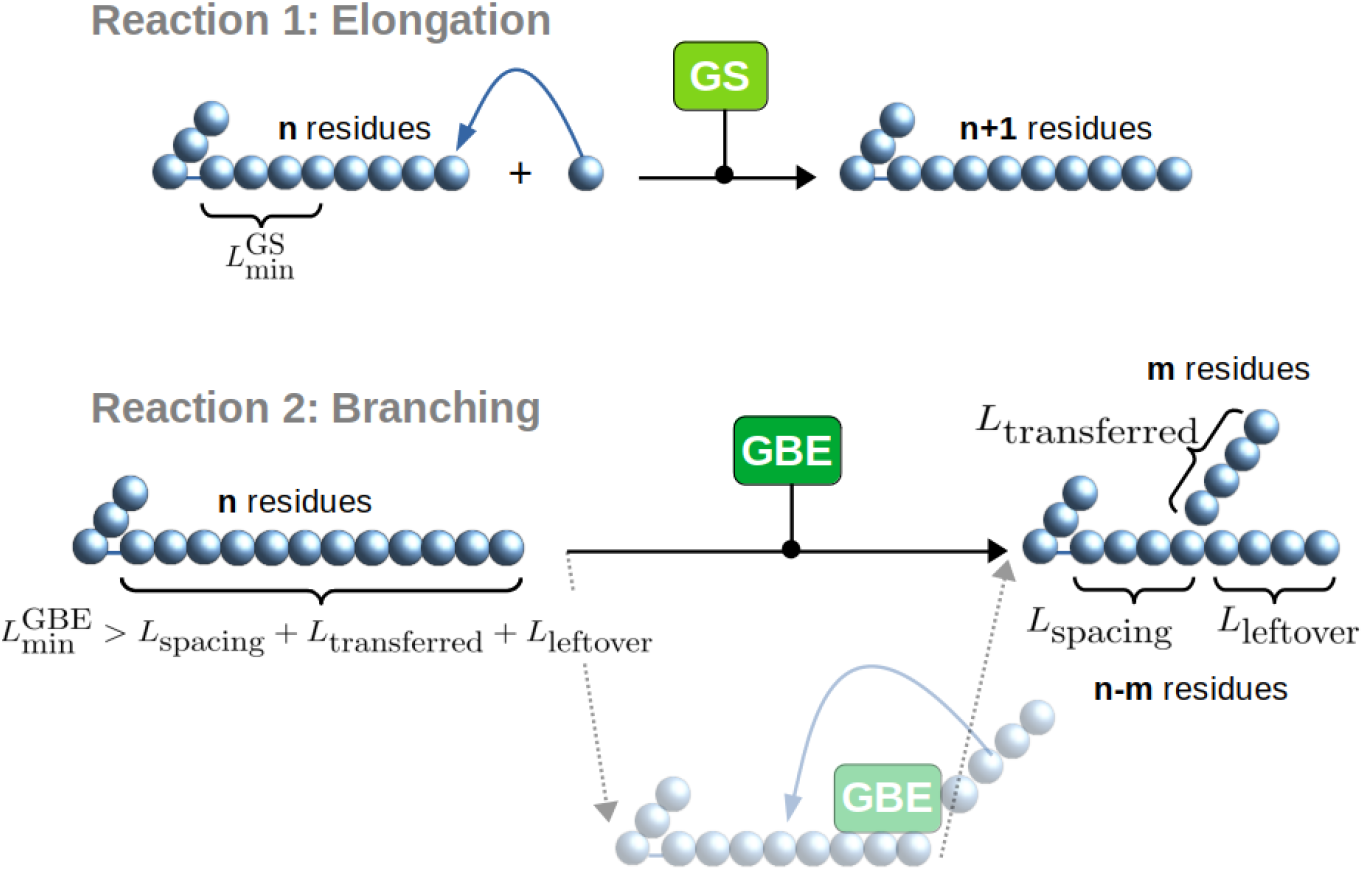
Glycogen synthesis reactions. Glycogen Synthase (GS) catalyses the elongation reaction. It needs a branch with a minimal DP as a substrate and a glucose unit to react. Glycogen Branching Enzyme (GBE) catalyses the branching reaction if the substrate’s DP is greater than the sum of 3 different minimal lengths.

GS binds the non-reducing end of an *α*-1,4 linear chain and elongates it by adding one glucose residue. It is commonly assumed that GS needs a glucose chain primer as a glucose acceptor [38]. In the model, we call 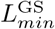 the minimal chain length of this required primer by GS. As elongation takes place, the chain becomes long enough to be cleaved and branched. This reaction is catalysed by GBE. Just like GS, the action of GBE is also characterised by specific substrate and product lengths as shown in Fig 3. Since less is known for GBE, we tested two models for its mechanism, namely the strict location model and the flexible location model. These are detailed in the Supplementary Material (see Figs 12 and 13). While comparing the two models to experimental data, we observed that the flexible location model provides a considerably better fit (see Figs 14 and 15 in the Supplementary Material). Thus, throughout the paper, we choose to use the flexible location model and will vary the GBE associated parameters. As illustrated in Fig 3, we consider a minimal chain length for the substrate (noted 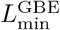) such that GBE is able to bind. We ensure that no daughter branch stands between the binding point and the non-reducing end of the branch. After binding, GBE cleaves at least 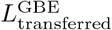 glucose units. Finally, GBE must attach the cleaved chain on the initial substrate, following an intramolecular process, and creating a new A chain. To precisely describe this last step, we define two additional lengths. First, the new *α*-1,6 branching point must not be closer than 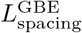 from either the first glucose of the chain, or an above *α*-1,6 branching point. Second, the new branching point must not be closer than 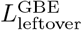 from the non-reducing end of the substrate chain, which is the original position of cleavage. Thus, for a new branch to be created by GBE, the substrate branch must have a chain length greater than 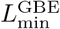, verifying:

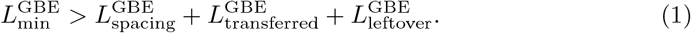

To illustrate the impact of these minimal lengths on the reaction outcomes, in Fig 4 we detail the case of 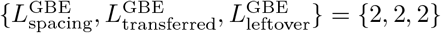 If the substrate reaches a length of DP equal 7, the condition (Eq 1) is fulfilled and the reaction may take place. If this reaction occurs, there is a single possible outcome (Fig 4, left panel). If the branching reaction occurs on a longer chain than just the minimal one, several outcomes are possible, all fulfilling the set of rules specified by the triplet {2, 2, 2}. Fig 4 (Right panel) depicts the case of a substrate chain with 9 residues, which results in 6 possible outcomes.

**Fig 4.**
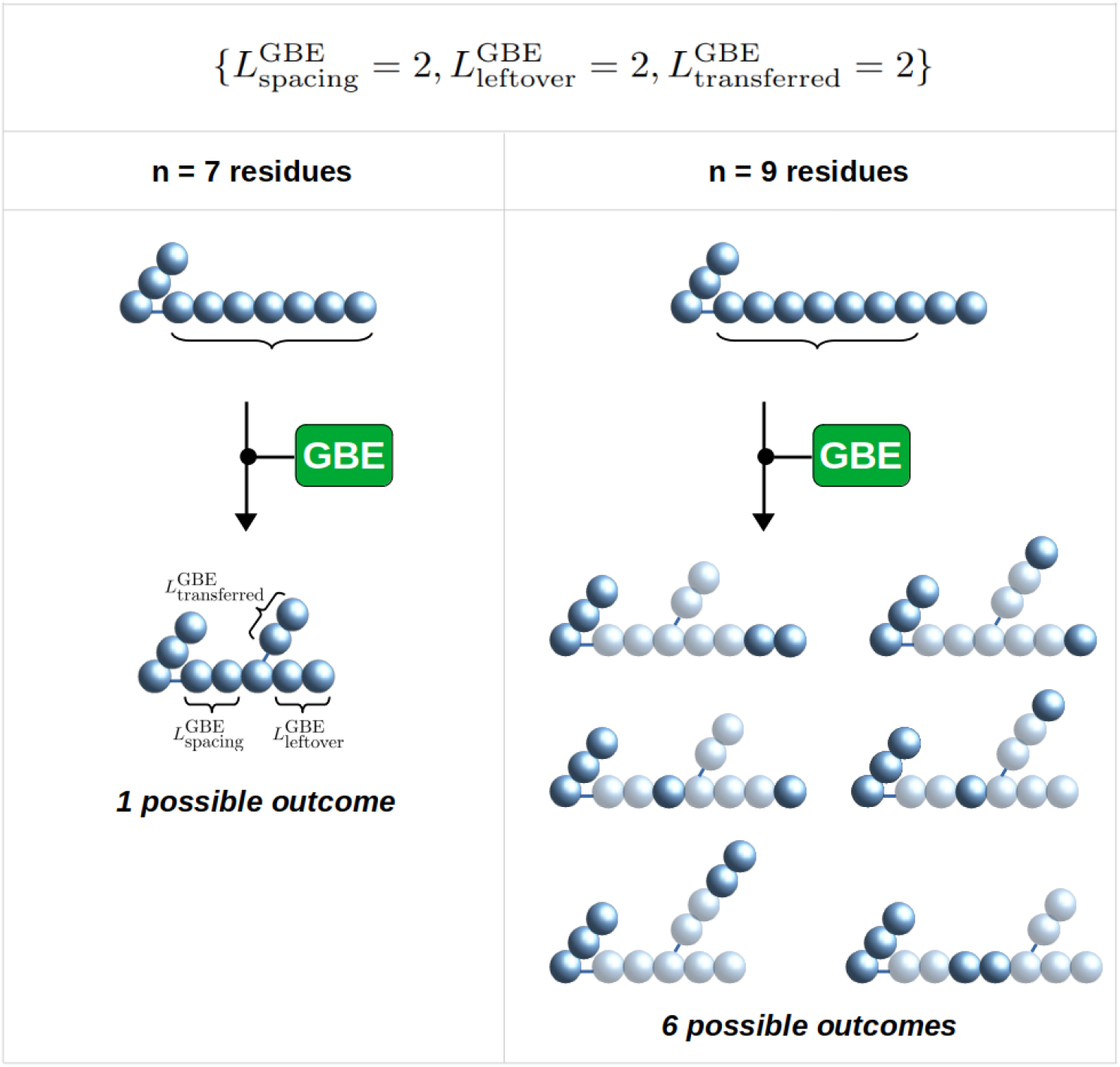
Illustration of the potential outcomes by GBE branching with 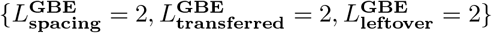. With these minimal lengths, the minimal DP required for a branching to occur is DP = 7. If the chain length is longer, the number of possible outcomes increases. **Left:** With a substrate of DP = 7, only one outcome is possible. **Right:** With a substrate of DP = 9, up to 6 distinct outcomes are possible.

### 2.2 Analysis of the model’s properties with generic parameters

#### 2.2.1 Elongation to branching ratio

When the simulation starts, the system is composed of a GN core with two primary chains standing in opposite directions in the center of the simulation space. Two enzymes (GS and GBE) modify the structure of the glycogen granule. To quantify that, we define Γ as the ratio of the GS over GBE reaction rates. For Γ ≈ 0, branching dominates over elongation, and *vice versa* when Γ ≫ 1. Fig 5 (Top) shows two simulated glycogen structures obtained with Γ = 0.2 and Γ = 10, respectively. We observe that a low Γ corresponds to a tightly packed structure, while a high Γ leads to a sparsely packed structure, with further elongated chains. Both simulations have been performed with a high *N* value, such that the number of monomers is not a limiting factor. The simulations end when the total number of monomers incorporated into the growing granule is *N* = 50,000. As can be seen in Fig 5 (Top), when Γ increases, for the same number of glucose units incorporated into the granule, the length of the chains increases while their number decreases, and so does the number of non-reducing ends (represented by green spheres).

**Fig 5.**
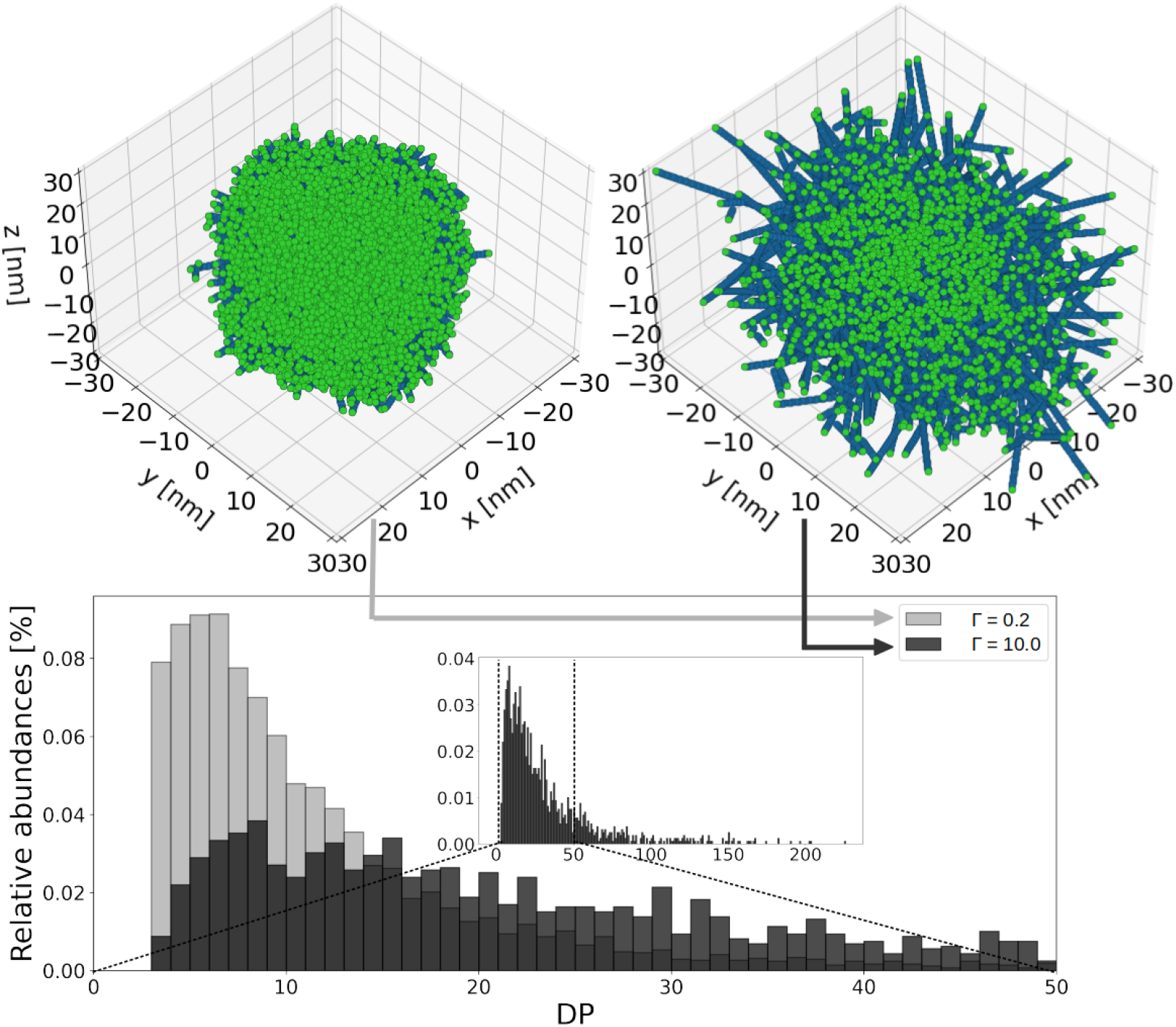
Simulated glycogen granule structures for different elongation to branching ratios (Γ). 50,000 glucose units are incorporated. **Top: 3D structures of glycogen granules.** Blue spheres represent the glucose units, green spheres the non-reducing ends. When Γ = 0.2, the structure of the granule is tightly packed. For Γ = 10.0, the structure of the granule is sparsely packed. **Bottom: Associated chain length distributions.** The light grey histogram shows the CLD for the tightly packed granule, while the black one shows that of the sparsely packed granule. The inset shows the full range of DP for Γ = 10.0. The longest chain is found to have a DP of 226.

Fig 5 (Bottom) shows the chain length distribution (CLD) for the two simulated structures. It is computed as the abundance of each chain length, taking into consideration all the chains of the structure. With Γ = 0.2 (light grey histogram) the average DP is 12.2 with a peak in the range [4 – 8] DP. When elongation is stronger than branching (Γ = 10.0, black histogram) the distribution shifts to higher DPs, the mean is 38.8, and the intensity of the peak is much reduced. By analogy with the well-studied case of starch, a chain length distribution with abundant high DP, might be an indication of double helix formation [39]. We do not model double helices as such, but our results allow to determine the Γ range that might lead to double helix formation, and thus potential glycogen precipitation.

#### 2.2.2 Granule density

Our approach tracks the position (*x, y, z*) of each glucose unit in three-dimensions. This allows us to compute how densely the granules are packed. Granule packing is quantified by the occupancy Ω, which is defined by the volume occupied by glucoses *V*_glucose_ divided by the total volume *V*_total_,

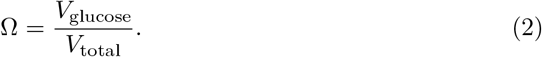

To characterise an entire granule, we consider the total volume *V*_total_ to be a sphere of gyration radius *R_g_* (defined in the section 5.1 of the Supplementary Material). To determine the occupancy at a given radius *r* from the center of the granule, we estimate the local occupancy in a spherical shell between radii *r* and *r* + Δ*r*. Fig 6 displays how the occupancy Ω as a function of the radius r dynamically changes during granule synthesis for two different values of the elongation to branching ratio Γ. The left panel shows the formation of a tightly packed granule (Γ = 0.2), while the right panel shows a sparsely packed structure (Γ = 10.0). Each line in the figure corresponds to the incorporation of 5,000 glucose units into the granule. It can be observed that granule synthesis proceeds in two phases. The first phase is characterised by an increase of the density close to the granule centre, while the radial extension increases only moderately. This can be explained by the fact that initially there is sufficient space to add new glucose units and there is hardly any steric hindrance among them. When steric hindrance constrains the synthesis (after around 10,000 glucose units have been incorporated), the system transits to the second phase. The latter is characterised by a radial expansion of the overall structure, while the density remains approximately constant around 30%. The two phases can be observed for both Γ values considered, but they are more pronounced for the tightly packed granule (Γ = 0.2).

**Fig 6.**
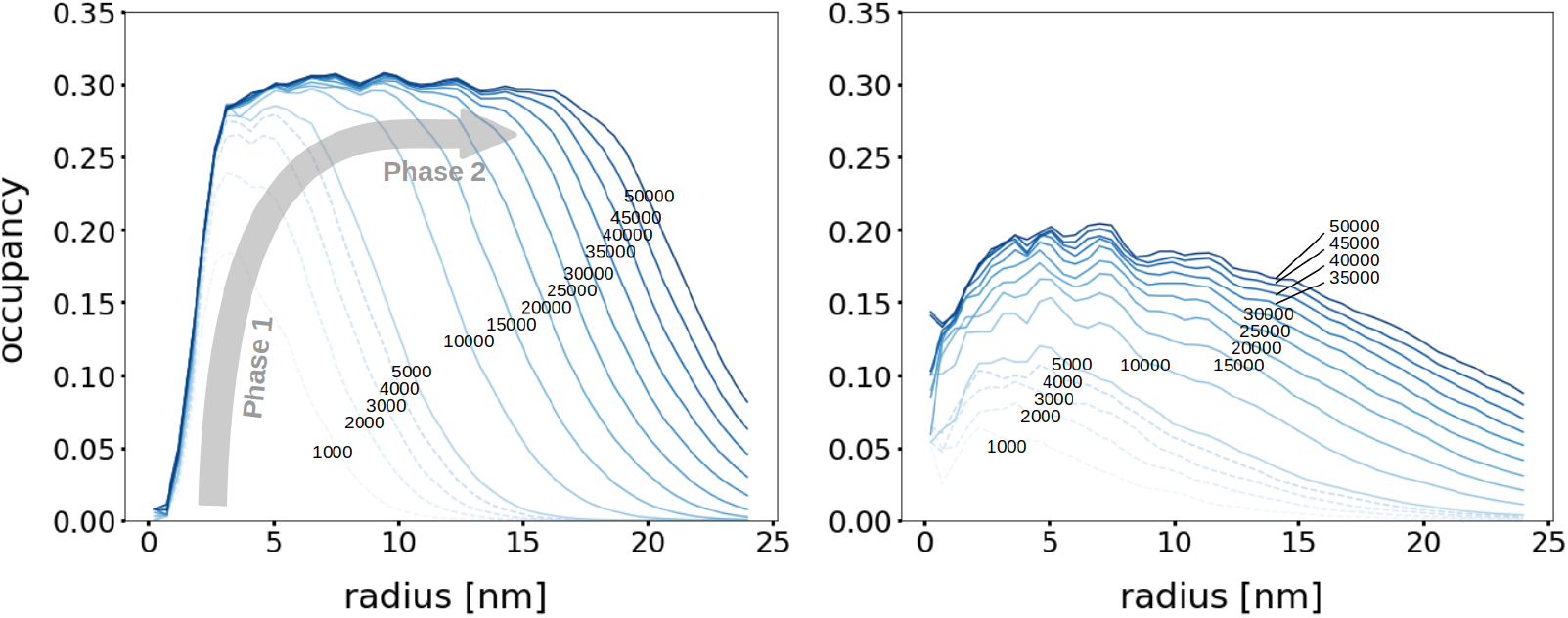
Dynamics of the occupancy profiles for a tightly (Γ = 0.2, left) and sparsely (Γ = 10.0, right) packed granule. Occupancy as a function of the radius at different simulation times. Each line corresponds to an incorporation of *N* = 5,000 glucose units. The simulation stops at *N* = 50,000. The grey arrow highlights the two phases of the granule synthesis dynamics. In phase 1, steric hindrance constrains are low, allowing occupancy to increase. In phase 2, i.e. after incorporation of ca. N = 10,000 glucose units, the occupancy reaches a plateau and the granule expands.

The relatively low occupancy close to the granule center is due to the presence of the GN core, which is not counted in the occupancy, but its corresponding volume cannot be filled with glucose units. We observe that at most 30% of the granule volume is occupied by glucose. This value is rather low, which is expected, since in the model, branches are straight helices, without any flexibility, while in reality, branches can bend and form locally higher densities. Specifically, this occupancy value is below Ω = 0.45, which we can estimate from previous studies [31]. There, the authors combine experimental and modelling approaches to conclude that granules of 55,000 glucoses have a radius of 21 nm [31]. Nonetheless, there are many uncertainties on the molecular masses experimentally measured. Thus, the occupancy value (i.e. Ω = 0.45) that we deduce from their work might hold large errors. For instance, it is unclear how the water and protein molecules embedded in the granules contribute to the molecular masses experimentally measured.

#### 2.2.3 Effect of the branching enzyme on the CLD

The branching enzyme mechanism is characterised by a triplet of integer numbers, denoted 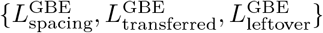, which specifies a unique set of rules for the enzymatic reaction. These rules considerably impact the CLD. Fig 7 shows that when these minimal lengths increase, the peak of the CLD is less pronounced and the distribution spreads towards higher DPs. Also, a change in each minimal length has a specific effect on the CLD.

**Fig 7.**
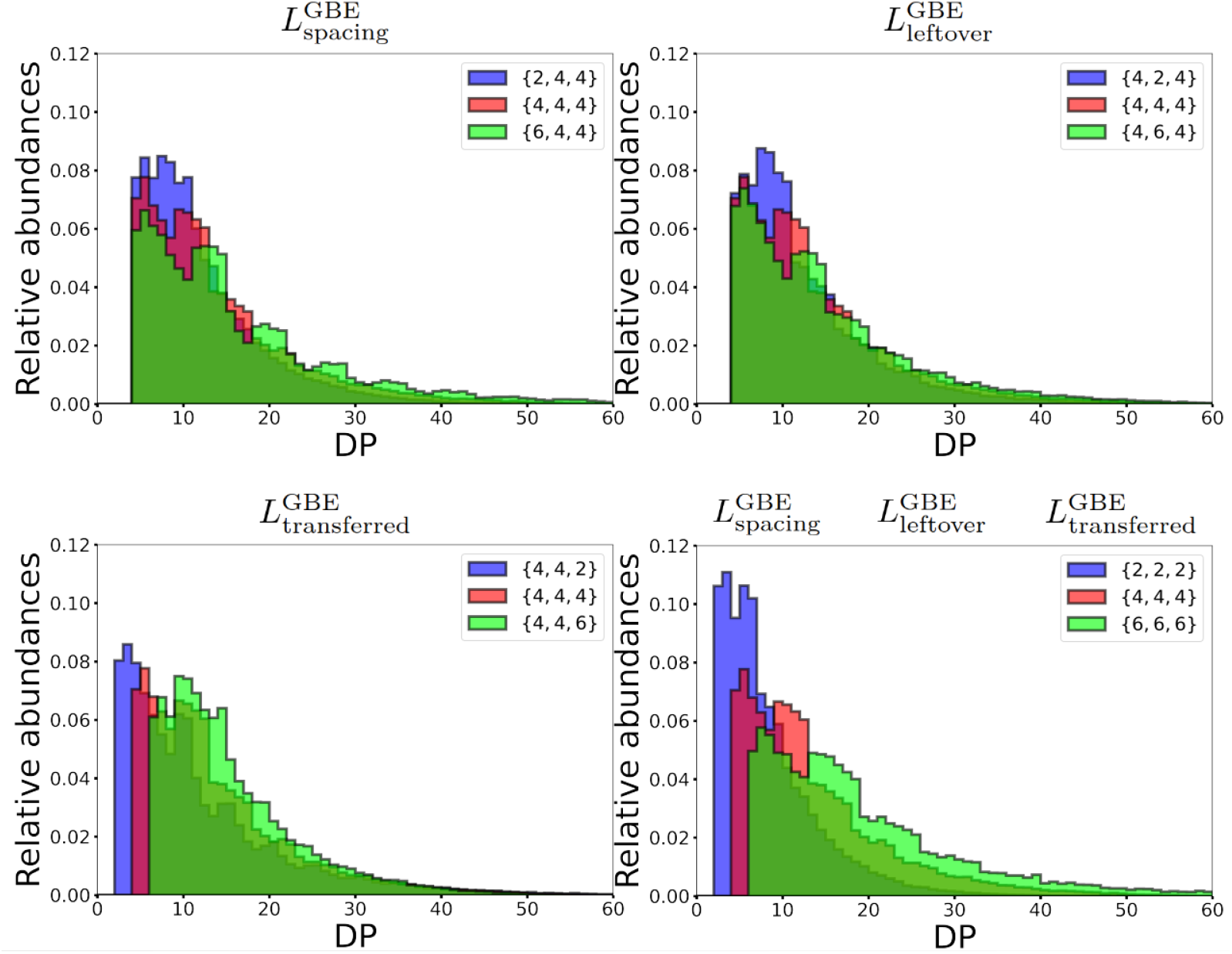
Effect of the chain length specificities on the CLD. **Top-left:** CLD for 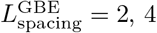, and 6, respectively. When 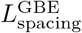 increases, a multi-modal distribution emerges. **Top-right:** CLD for 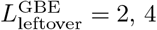, and 6, respectively. When 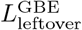 increases, the peak is reduced and the overall distribution spreads towards higher DPs. **Bottom-left:** CLD for 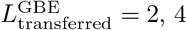, and 6, respectively. When 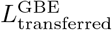 increases, the distribution shifts towards higher DPs. **Bottom-right:** CLD 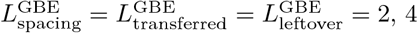, and 6, respectively. Varying these distinct minimal lengths concomitantly, combines the individual effects described above, when a single length is varied. Each CLD is the result of averaging 200 simulations of granules with 5,000 glucose units each.

Increasing 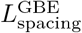 drastically reduces the peak and spreads the distribution, while making it bimodal. An increase in 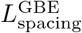 reduces the granule’s number of possible configurations. Less configurations are possible, and 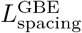 further shapes the chain length distribution. Chains of DP that are combinations of 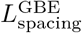 and 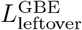 are favoured, resulting in local peaks. Increasing 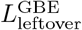 also decreases the peak and spreads the overall distribution towards higher DPs. Besides, 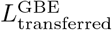 has a different effect on the structure. Since 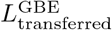 is the smallest DP that can be formed, it is found on the leftmost part of the CLD, and variations in 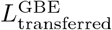 shift the overall distribution by the corresponding exact amount.

It is important to notice, that these results are obtained when branching dominates over elongation. Increasing the elongation to branching ratio Γ systematically smoothens any multi-modal CLD, because it introduces flexibility in the branching location. It also flattens the peak and spreads the distribution towards higher DPs. Consequently, bi-modal distributions are obtained for high 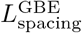 and low Γ values.

As discussed in the Elongation to branching ratio section, we can compare the synthesis process of glycogen and starch. Specifically, in starch the CLD is bi-modal and shifted towards higher DPs as compared to glycogen [40]. Based on our preceding remarks, it could mean that starch branching enzymes are characterised by large substrate specificity lengths, corresponding to a more constrained mechanism than for glycogen. An alternative explanation for the arising of multi-modal CLDs, also based on highly constrained branching, is discussed in the Supplementary Material (see Figs 12 and 13).

### 2.3 Comparison to experimental data

#### 2.3.1 Parameter calibration

Based on our simulations, it is clear that the CLD is a signature of the branching enzyme mechanistics and substrate specificity. This signature results from the combination of many different parameters. Therefore, it is particularly difficult to analyse experimental data and infer parameter values without a complementary modelling approach. To extract useful information, and compare simulations to experimental data, we designed a fitting procedure, which is detailed in the Material and Methods section. With this strategy, we are able to determine parameter values that we compare to experimental results.

GBE’s mechanism and substrate specificity are incompletely characterised, yet they drastically impact on the CLD, as shown in Fig 7. Therefore, we specifically focus on this enzyme. To this end, we use experimental data obtained by Sullivan et al. [23] for mice muscles, that we extracted from their publication using the software Engauge Digitizer [41]. After purification, the granule chains are debranched using isoamylase, and their degree of polymerisation is measured using size exclusion chromatography. Our fitting procedure can be applied to any glycogen data, obtained from any specie and tissue. The data by Sullivan et al. [23] present two major advantages for our study. First, they are quantitative measurements of good resolution. Second, the authors investigated the case of a defective GBE.

Fig 8 shows the heat-map containing the best fit we obtained with the model. The minimal lengths for the branching enzyme are fixed to 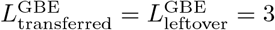, while 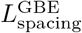 is represented on the Y-axis, ranging from 1 to 6. The elongation to branching ratio Γ is varied on the X-axis. We notice that only low values for 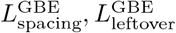 and 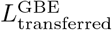 allow to fit the experimental data. However, one should remember that these are minimal lengths and that the positions at which GBE is able to cleave and branch is flexible beyond these minimal lengths (see Fig 4). Overall, not only have we ran our fitting procedure using both the flexible and the strict location branching models, but also considering distinct values for *ρ* (see Figs 14 and 15 in the Supplementary Material). Interestingly, the best fit is obtained for *ρ* = 0.65 nm, which reflects the realistic size of a glucose unit inside an helical chain. This highlights the role of the steric hindrance to mimic the granule’s structural properties. The best fit is obtained for the following set of parameters:

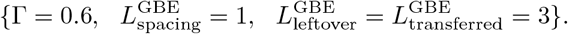

**Fig 8.**
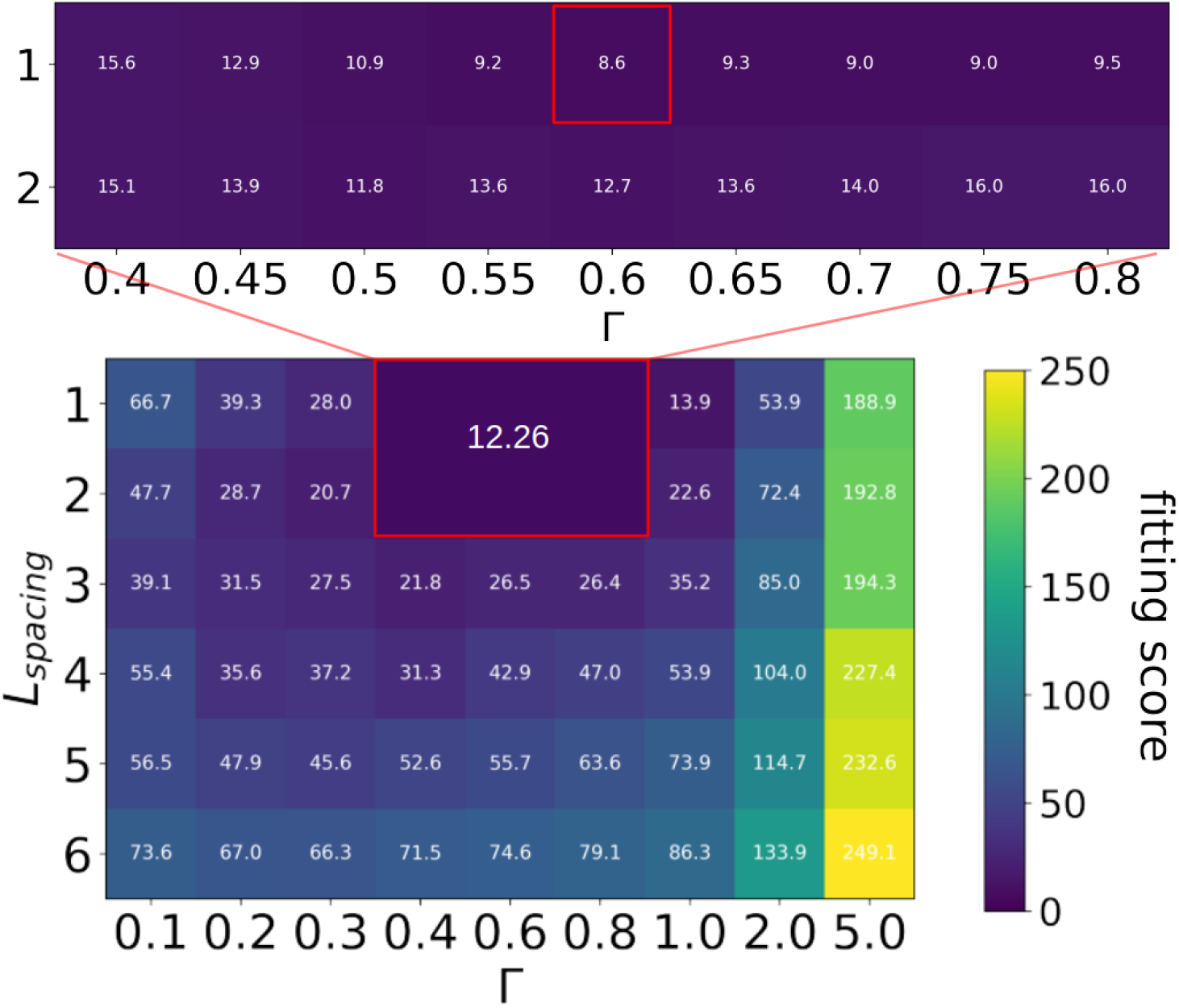
Heat-map showing fitting scores for various sets of parameters. The *Y*-axis shows 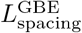 ranging from 1 to 6. The *X*-axis shows the elongation to branching ratio Γ ranging from 0.1 to 5.0. A given cell corresponds to a parameter set 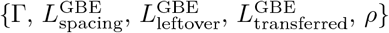. Additional parameter sets are tested around good scores, i.e. the resolution on the elongation to branching ratio is increased, as well as the number of runs averaged. This area is surrounded by a red rectangle in which the average score is 12.26. Fitting scores are ranging from 8.6 to 369.0. The best score is 8.6 (red square in the inset heat-map) which corresponds to 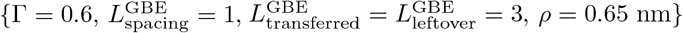.

Importantly, the parameter values for the branching enzyme inferred from the best fit are distinct from those reported in the field [31, 42, 43], especially for the typical spacing observed between two branches [31]. Also, based on our results, GBE is able to transfer less than 4 glucose units. Knowing that GS’s chain length specificity has to follow the same rule, this questions the commonly assumed value of DP4 as the minimal length that can be elongated by GS.

In Fig 9, the CLD for the best fit is shown, together with simulation results for parameter values typically reported in the experimental literature, i.e. 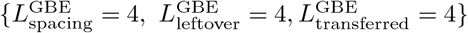. For the latter, we observe a plateau from DP4 to DP10, while experimental data show a peak between DP6 and DP8. Additionally, longer chains (DP ≥ 15) are over-represented in our results. Noticeably, for this set of GBE minimal lengths, our model is not able to reproduce the experimental data by Sullivan and coworkers [23], even when varying the elongation to branching ratio Γ (see Fig 14 in the Supplementary Material).

**Fig 9.**
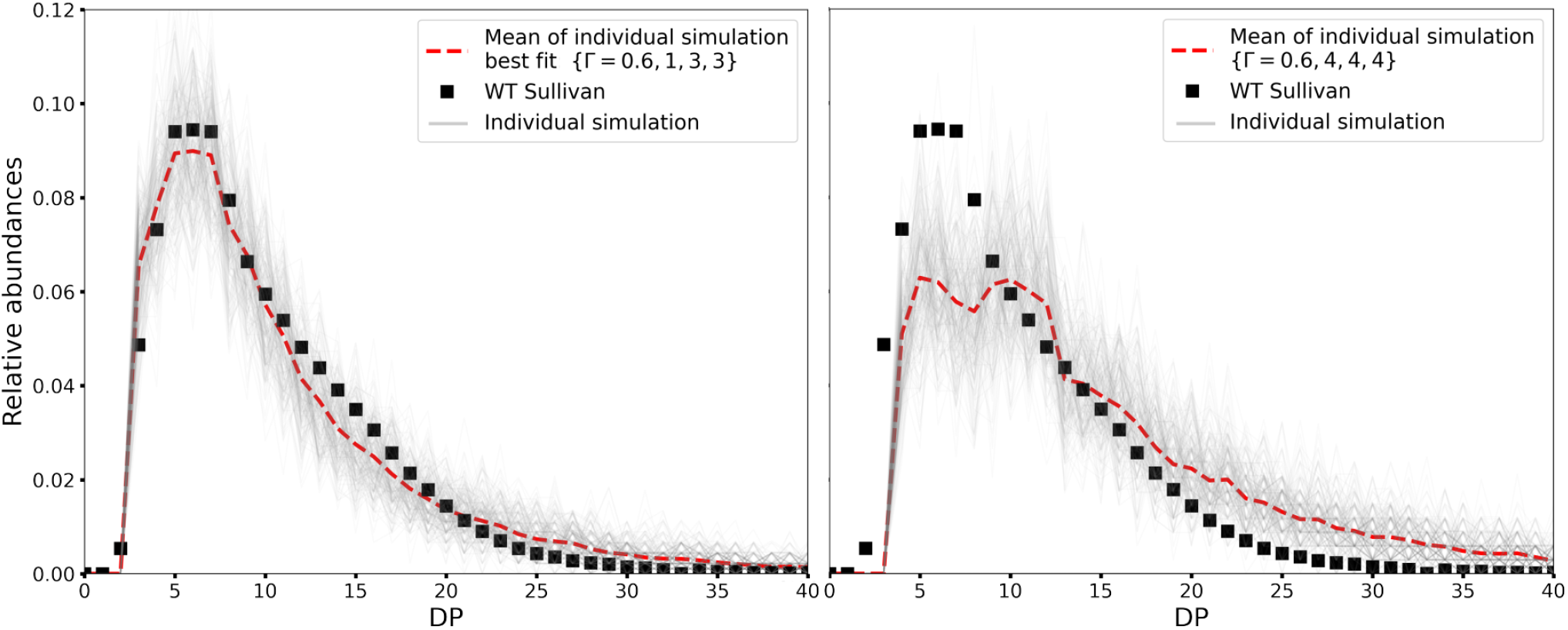
Comparison of simulated *versus* experimental CLDs. Experimental data are from Sullivan and coworkers [23] (black squares). In each simulation run 50,000 glucose units are incorporated in the growing granule (grey line). The average over 200 runs is represented as a red dotted line. Left: The CLD for the best fitting score 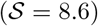 is obtained with 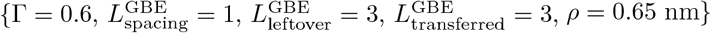. Our best fit almost perfectly captures the experimental CLD. Right: CLD using parameter values typically assumed in the literature 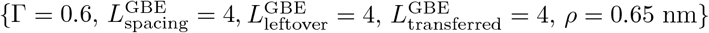 [42, 43]. The simulated CLD differs a lot from the experimental CLD, with under-representation of small DPs, and over-representation of high DPs.

#### 2.3.2 Glycogen structure using the fitted parameters

In this section, unless otherwise specified, we assume that GBE chain length specificities are set to those of the best fit 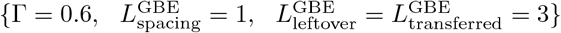. With these Parameters, we simulate the synthesis of glycogen granules, and compute their structural features and macroscopic characteristics (see Table 1). For each of those, the average values and standard errors are calculated over 10 granules with *N* = 50,000 glucose units. The number of non-reducing ends (noted *N*_NREs_) is equal to the total number of chains, since there is one non-reducing end per chain.

**Table 1.** Summary of the granule structural features for the best fit parameters.

|  | Structural features |  |  |  |  | Macroscopic characteristics |  |
| --- | --- | --- | --- | --- | --- | --- | --- |
| | $N_{\text{NREs}}$ | A:B | CL | Branching degree | Generation | $\Omega$ | $R_g$ |
| $\Gamma = 0.6$ | $4136.0 \pm 39.7$ | $0.98 \pm 0.02$ | $12.09 \pm 0.12$ | $0.0902 \pm 0.0009$ | $22.1 \pm 0.9$ | $0.504 \pm 0.012$ | $21.95 \text{ nm} \pm 0.16$ |
| $\Gamma = 50.0$ | $424.8 \pm 23.9$ | $1.03 \pm 0.08$ | $118.08 \pm 6.74$ | $0.0085 \pm 0.0004$ | $10.6 \pm 1.4$ | $0.026 \pm 0.004$ | $52.36 \text{ nm} \pm 2.91$ |

For Γ = 0.6, we find that on average, granules are made of 4,136 chains, with a chain length of 12.09 glucoses. These two quantities are intrinsically linked since the total number of glucoses is fixed to *N* = 50,000. The higher the average chain length (noted CL), the lower the total number of non-reducing ends, since CL · *N*_NREs_ = N. The A:B ratio is found to be 0.98. With *N*_NREs_ = 4,136, it means that ca. 2,047 chains are of type A, while 2,089 are of type B. The branching degree is found to be 9.02%. This is in good agreement with the range typically reported in glycogen (i.e. [5-10 %] [9, 27]), while starch exhibits much lower values (typically between 1 and 5% [24–26]). The average occupancy is Ω = 0.504, which means that half of the total granule volume is filled with glucoses. The gyration radius is *R_g_* = 21.95 nm which is consistent with radii reported for big granules (molecular weight ca. 10^7^g.mol^−1^), like the ones considered here (N = 50,000) [13, 44].

It is interesting to notice that ca. 21 nm is the upper limit for the radius of a glycogen granule of ca. 50,000 glucoses, when considering the fractal structure depicted by Meléndez et al. [6]. Thus, with our approach, that is not based on a fractal structure, we determine a gyration radius close to their results. However, the glycogen structures we simulate are deeply distinct from theirs. In our simulations, we observe that the density is approximately constant with the radius, while a fractal glycogen granule has a density that increases exponentially with the radius, so resembling a rather empty shell. Also, we show that our simulations can reproduce characteristic quantities (summarised in Table 1 for Γ = 0.6), which are in good agreement with experimental results. Thus, our model appears to provide a more realistic depiction of glycogen granules.

For comparison, in Table 1, we also report the case of Γ = 50.0, while keeping all other parameters unchanged. By increasing Γ, we force elongation over branching. As expected, less chains are created but they are longer. Consistently, the branching degree is also lower. In our model, chains are rigid rods that cannot bend, which leads to a bigger radius of gyration. Since the total content of glucose is the same like for Γ = 0.6, the occupancy Ω decreases almost proportionally. *In vivo*, glycogen granules might follow a distinct behaviour, where the presence of long chains could trigger precipitation, like for instance reported in the Lafora and GBE diseases [23, 45].

The formation of an A chain results from branching either an A or a B chain. These different events can be represented as reactions and described as follows:

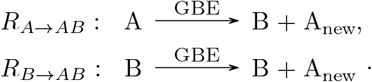

To illustrate how different branch structures correspond to specific A:B ratios, we sketch extreme cases in Fig 10. If the *R_B→AB_* and *R_A→AB_* reactions are equally likely, a purely probabilistic approach predicts that, on average, the A:B ratio is equal to 1, independently of the relative activities of the enzymes (see Fig 16 in the Supplementary Material). Such a ratio is for instance observed in the first two rows of Table 1, with A:B = 0.98 ± 0.02 and A:B = 1.03 ± 0.08, which is in good agreement with reported values for glycogen (A:B = 0.6 – 1.2 [29]). Other closely related branched polysaccharides can exhibit different A:B ratios. For example, amylopectin has a typical A:B ratio in the range from 1.5 to 2.6 [29, 30], depending on the organism. It is therefore interesting to investigate which factors influence this ratio.

**Fig 10.**
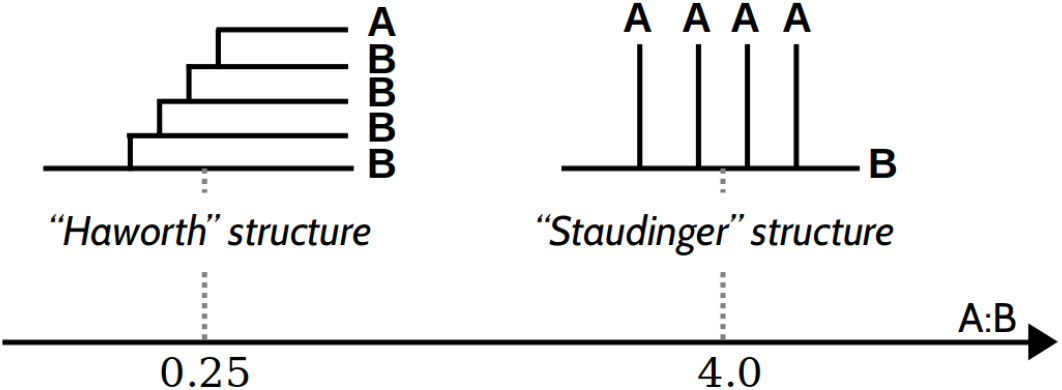
Two schematic branched structures with different A:B ratios. For a “Haworth”-like structure, branching reactions tend to occur on A chains, leading to a low A:B ratio, while for a “Staudinger”-like structure branching on B chains is favoured, leading to a high A:B ratio.

Here, we identify two effects of the interplay between the branching mechanism and the dynamics of the granule structure, highlighting once again the importance of the branching enzyme on the emerging structural patterns. First, considering a small Γ, branching dominates over elongation, and the enzyme branches as soon as it can. Thus, if in addition 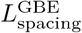 is small, the A chains are closely stacked together and the granule is tightly packed. In this regime, overall, A chains are closer to other A chains than to B chains. Due to the increased steric hindrance for A chains, B chains react more easily, which favours the *R_B→_AB* reaction, and leaves A chains unchanged. Therefore, the A:B ratio increases, meaning that it shifts to the right along the axis in Fig 10. Second, if 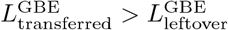, when a branching occurs, on either an A or a B chain, the newly formed A chain will on average be longer than the B (leftover) one. Therefore, the new A chain will react faster, favouring *R_A→AB_*, and lead to a reduction of the A:B ratio, meaning that it shifts to the left along the axis in Fig 10. We confirmed this effect for the specific case of 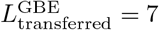 and 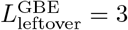. The resulting structures have an A:B ratio on average equal to 0.80 ± 0.02. If the values of the minimal lengths are exchanged, such that 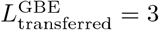 and 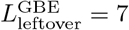, the A:B ratio is instead equal to 2.01 ± 0.04. It appears that the 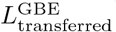 and 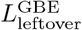 are specific mechanistic features that uniquely determine the A:B ratio. Consequently, they could cause the main structural differences observed between amylopectin (the prime constituent of starch) and glycogen.

## 3 Material and Methods

### 3.1 Simulation procedure

We present a model that simulates the dynamics of glycogen synthesis. We record all enzymatic reaction steps, the time at which they occur, and the full 3-dimensional details of the glycogen structure at any time point. The model specifically keeps track of each glucose position, the complete connectivity of the chains, and the position of each branching point. To account for this complexity, we implement the model using a stochastic algorithm. This approach also allows to specifically consider how changes in the glycogen structure enable or disable enzymatic reactions. For comparison, employing a deterministic approach, for instance based on systems of ordinary differential equations, would lead to unnecessarily complex simulation rules, that would also include additional ad-hoc assumptions. Besides, randomness has a stronger impact on a system as it involves small numbers, like the one we consider here. Indeed, the synthesis and breakdown of a single glycogen granule of ca. 50,000 glucose units involves only a small number of enzymes. For example, experiments indicate that on average a single glycogen synthase enzyme is active per granule [46]. As a result, a stochastic approach appears very natural to deal with this complex and spatially structured substrate.

We base our stochastic approach on the Gillespie algorithm [47]. At each time step, it consists in randomly selecting both an enzymatic reaction and its duration, while systematically updating the structure of the glycogen molecule accordingly. While the enzymatic reaction is selected from a uniform distribution, the time is chosen from an exponential distribution, such that the Gillespie algorithm allows to simulate the real time of the dynamics of the system, as far as the underlying enzymatic mechanisms and their kinetic parameters are known. Each reaction is associated to a specific propensity, that we choose as proportional to the concentration of enzymes and substrate, thereby following a typical mass-action kinetics.

The Gillespie algorithm accounts for all possible reactions, and keeps track of any modification of the granule structure, such that only possible reactions can be selected. Nonetheless, because of the complexity of the structure we simulate, to account for steric hindrance among monomers, we instead allow for certain reactions to be rejected. For these rejected reactions the time elapsed is not accounted.

### 3.2 Best fit algorithm

The model contains various kinds of parameters. Some describe the physical properties of the glycogen structure, others relate to the enzymatic activity, including the enzyme substrate specificities. On the one hand, certain parameters are inferred from literature data, for instance the minimum DP for GS to act 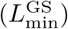. On the other hand, other parameters are free to be fitted by our in-house designed best fit algorithm. Here we choose to fit the minimum DP between two branches after a branching reaction 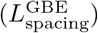, the minimum DP between the non-reducing end of the mother branch and the new branching point 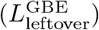, and the ratio between the elongation and branching reaction rates (Γ).

The best fit algorithm consists in setting bounding ranges for the parameters to be fitted, and systematically scanning the parameter space thereby defined. For each parameter set tested, we run 50 simulations, take the average of the resulting CLDs, and compare it to our targeted experimental data set. For each parameter set, the comparison to the experimental data is measured by the score 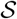, defined as follows:

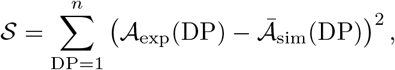

where 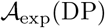 is the experimental CLD abundance for a given DP and 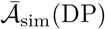 the average simulated CLD abundance for a given DP. Therefore, 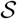 measures the difference between the average simulated and the experimental CLDs. The best fit is found for the parameter set that has the lowest score 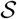.

## 4 Conclusion

### 4.1 Discussion of the results

According to the World Health Organization, metabolic diseases are a burden in western countries. Among them, Glycogen Storage diseases (GSDs), Lafora disease, Adult Polyglucosan Body Disease (APBD), and even diabetes directly or indirectly involve glycogen. Investigating the regulation of glycogen’s structural properties could therefore strongly contribute to further understand such diseases. Computational models that encompass the complex interplay of both glycogen’s structure and its metabolism allow to tackle this challenge but remain poorly exploited.

In this article, we present a stochastic structural model for glycogen synthesis. The model provides a precise description of both the structure of glycogen in 3 dimensions, that we can visualise, and the detailed dynamics and mechanistics of the underlying enzymatic reactions. For instance, both elongation and branching have precise rules regarding their substrate specificities. Modelling glycogen granules in 3 dimensions is made possible by a coarse-grained geometrical description at the glucose level, allowing us to track all the glucose units in space. This description also accounts for the steric hindrance effects resulting from the impossibility for the chains to overlap. Simulating reaction dynamics relies on a Gillespie algorithm, which determines when and where a reaction of branching or elongation takes place, thereby dictating the corresponding change in the 3-dimensional structure.

We show qualitatively how enzyme activity affects glycogen structure for generic sets of parameters. We highlight two different synthesis regimes, depending on Γ, the ratio between the elongation and branching reaction rates. By varying Γ, either small and dense, or big and sparse granules can be simulated. Though, considering chains bending and intermolecular interactions might lead to more complex results, in particular during the synthesis of sparse granules. The phenomenology of the two synthesis regimes is also confirmed by the occupancy profiles along the radius of the corresponding granules. In both synthesis regimes, first the center of the granule is filled around GN, before reaching a critical density preventing further internal reactions to occur due to steric hindrance. Then, the granule expands such that the density remains approximately constant within the sphere defined by the gyration radius. While this result is a consequence of the geometrical assumptions of our model, it is interesting to note that a glycogen fractal description would instead give a density exponentially increasing with the granule radius. Besides our own results and the various arguments exhibited against a fractal representation of glycogen, we would like to highlight here that, as soon as one considers even a single degree of freedom in the glycogen branching reaction, it would rapidly lead to the loss of any “fractal-like” structure. Such degrees of freedom are necessarily present in natural conditions. For instance, the dihedral angles defining the *α* – 1, 6 bonds may take various values, making a fractal pattern very unlikely *in vivo*. Beyond these spatial considerations, the model establishes a natural and clear connection between enzymes’ reaction rates, and both the degree of polymerisation of the chains, and the number of non-reducing ends.

Our results show that the chain length distribution (CLD) of glycogen is highly sensitive to the branching reaction, predominantly its mechanism. Each of the three characteristic minimal lengths of the reaction has specific effects on the CLD. Additionally, if any of them increases, less branching outcomes are possible, eventually leading to a bi-modal or even multi-modal distribution. This effect is enhanced by high branching activity. When varied together, these minimal lengths show even more complex imprints on the CLD. In contrast, multi-modal distributions become less pronounced by increasing the elongation reaction rate, because rapid elongation increases the number of possible configuration outcomes. Additionally, increasing elongation leads to longer chains and results in a CLD spreading towards higher DPs. Altogether, we show that the CLD, and in particular peaks location and intensity, are subtly affected by several complex effects.

Guided by these findings, we propose to consider the CLD not only as an important structural feature of glycogen, but also as a signature of GBE. Thus, fitting experimental data with our model arises as a natural strategy to infer knowledge on the GBE mechanism. Not only did we illustrate the strength of our fitting procedure on an experimental data set from Sullivan et al. [23], but we also extracted several macroscopic characteristic features from the resulting fit, which we compared to various literature sources, thereby confirming the validity of our results. Using this fit, at the microscopic level, we were able to discriminate between the two branching models we hypothesised, and selected the flexible location branching over the strict one. Besides, we could critically evaluate parameter values typically reported in the literature. For instance, it is often assumed that GBE transfers branches of DP 4 or 6, or even longer [31, 42, 43]. Similarly, it is typically reported that around 6 glucose units space two branches. Within the framework of our model and its underlying assumptions, it is impossible to reproduce CLDs of *in vivo* glycogen using the above mentioned values. Instead, our fitting procedure suggests that a high flexibility is necessary for both the branching mechanism and the minimal lengths involved. This finding fully confirms the importance of modelling glycogen synthesis using a stochastic approach.

Moreover, our customisable branching model shows that the A:B ratio is independent of the kinetic parameter Γ, but specifically determined by the difference between 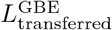 and 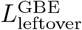. Based on these observations, we hypothesise that the branching mechanism is chiefly responsible for the structural differences observed between starch and glycogen.

Throughout this study, our coarse-grained approach accounts for the contribution of individual glucoses to the overall granule structure, by considering them as spheres of radius 0.65 nm. It is interesting to notice that if we set the glucose volume to 0 nm^3^, the CLD remains almost unchanged while other macroscopic properties of the granule, such as its overall volume, are dramatically impacted. This observation confirms once more that the glycogen CLD is primarily shaped by the enzyme mechanisms.

### 4.2 Outlook

It is important to keep in mind that our model contains limitations that may be circumvented by further refining the model assumptions. In our model, the simplified coarse-grained representation of glucoses assumes that all of them are arranged in single helices. This hypothesis implies that all *α* – 1, 4 glycosidic bonds have the same angle values. Yet, *in vivo*, this is highly unlikely, and instead, the dihedral angles of the *α* – 1,4 bonds should be able to take various values. To take this into account, we could randomly pick the dihedral angles of the *α* – 1,4 bonds using the Ramachandran plots of their energetically favourable regions. As a first trial, we could use that of maltose, that is well characterised [48]. We expect that, introducing such disorder in the angles, chains will appear longer. Besides, in the model, we consider that all chains are stiff. Thus, to improve the macroscopic representation of the chains, we could introduce the possibility that they bend when encountering steric hindrance. To do so, we would minimise their torsion energy, like it is done in polymer physics models [49]. Opposite to the change suggested for the *α* – 1,4 bonds variability, accounting for the flexibility of the chains might lead to a higher granule density, and thus, a lower radius. The fact that these two effects might cancel each other, possibly explains why our simplified model is nonetheless able to capture realistic *in vivo* granule radii. Besides, in abnormal conditions, the potential formation of double helices may not only lead to glycogen precipitation, but also prevent enzymatic reactions. Thus, a later improvement of the model could include to tune the enzymes’ reaction rate depending on the substrate chain configuration and length.

Throughout this article, we focus on glycogen synthesis, yet, simulating the degradation dynamics with the algorithms we developed would be straightforward. We expect residual degradation activity to only lightly modify the effective elongation to branching ratio Γ, and slow down the synthesis. In such a case, the CLD would be slightly shifted to the left, and the first chain length detected would be the smallest value between 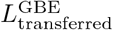 and its GP counterpart. It could be particularly interesting to account for the degrading enzymes, beyond a residual activity, in case the synthesis is defective. Indeed, abnormal structures produced by defective synthesis enzymes could be degraded, and thereby corrected, by degrading enzymes. For instance, if 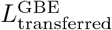 becomes too short for GS to elongate the newly formed chain, GDE could unbranch the latter and thereby preserve the macroscopic properties of the granule. Last, following an approach analogous to the one taken in this article, one could choose to investigate the glycogen granules’ breakdown in full depth, by first synthesising granules and then proceeding to their degradation. Although, for the sake of simplicity, we would then uncouple in time the synthesis and degradation processes, it would still be very interesting to study the mutual impact of distinct modes of synthesis and degradation on the overall glucose release and fixation.

In this article, we show that the availability of the substrate strongly influences the enzyme activity, leading to distinct chain lengths and number of non-reducing ends. Noticeably, other modelling approaches, such as metabolic ODE models, instead consider glycogen as a single metabolite approximated by the sum of all glucose units that compose it. A direct consequence is that any structural aspects are neglected and these models cannot differentiate between a single chain of 50,000 glucoses, and an actual granule of the same weight. Still, for instance, the number of non-reducing ends available for elongation are drastically different in these two cases. Thus, coupling our model to glycogen metabolic ODE models, would constitute a hybrid approach that would include key structural details, while enlarging its biochemical scope. It would thereby open up a whole new range of modelling possibilities. For instance, it would allow to investigate the interplay between glycogen structure and the evolution in time of important metabolites, under various physiological conditions, including diseases scenarios. Using this approach, we shall be able to characterise the phenomenology of each glycogen storage disease, with a focus on the role of glycogen structure, and address questions such as glycogen accumulation, glucose cycling, glucose homeostasis, and even glycogen precipitation.

## Data availability statement

The source code and data used to produce the results and analyses presented in this manuscript are available on a GitHub repository at https://github.com/yvanrousset/Stochastic-modeling-of-a-three-dimensional-glycogen-granule

## Acknowledgments

This paper is supported by European Union’s Horizon 2020 research and innovation program under the Marie Skłodowska-Curie grant agreement PoLiMeR, No 812616. O.E. and A.R. are supported by the Deutsche Forschungsgemeinschaft (DFG) under Germany’s Excellence Strategy EXC 2048/1, Project ID: 390686111. The current position of A.R. is funded by the German federal and state programme Professorinnenprogramm III for female scientists.

## 5 Supplementary materials

### 5.1 Symbols definition

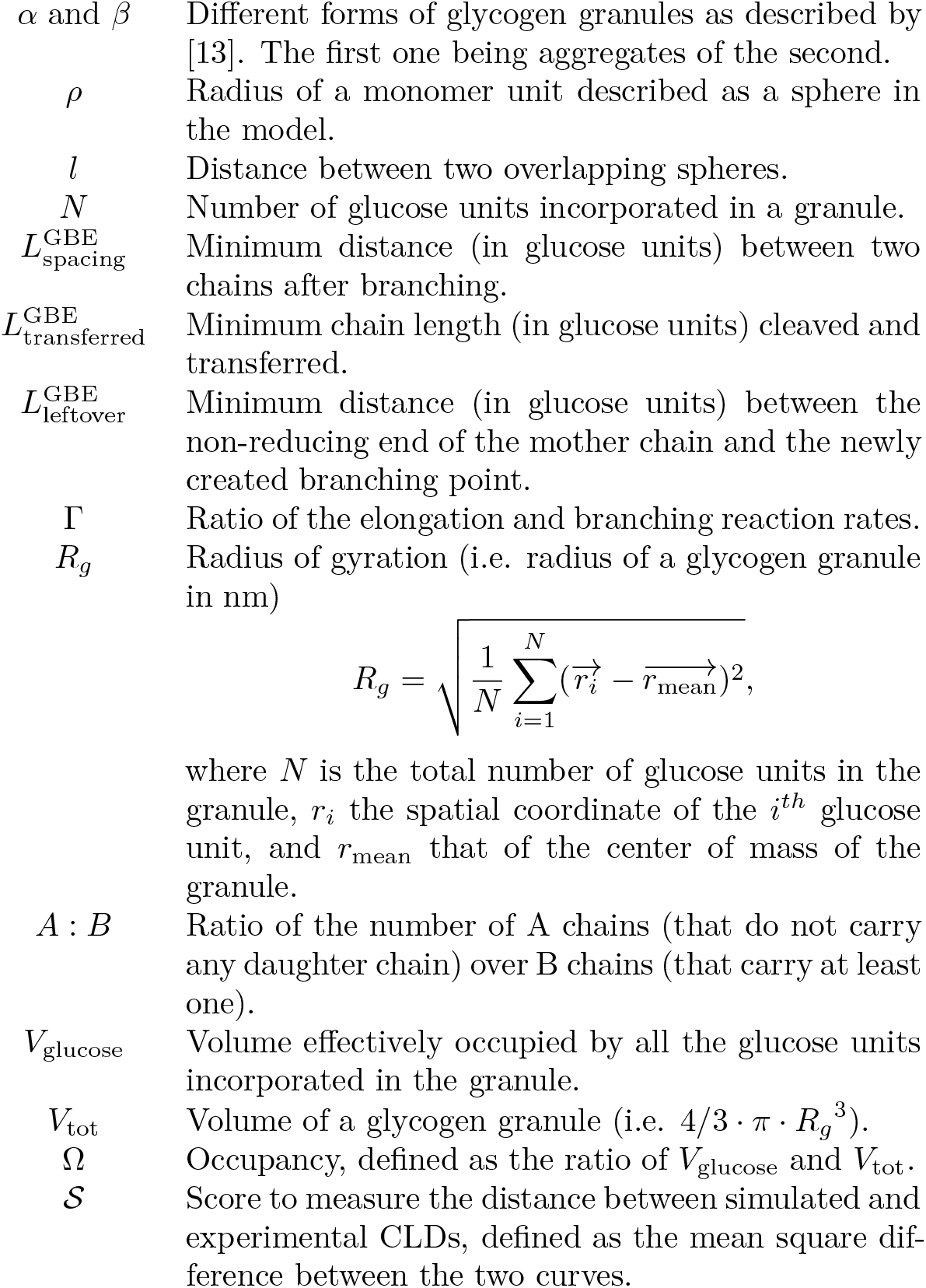

### 5.2

#### Abbreviations list

GS: Glycogen synthase
GN: Glycogenin
GBE: Glycogen branching enzyme
NREs: Number of non-reducing ends
DP: Degree of polymerisation
A:B: Ratio between the number of A chains and the number of B chains
CL: Chain length

### 5.3 Effect of self-exclusion

Modelling the 3D structure of glycogen, while considering self-exclusion of glucoses, allows investigating glycogen chain patterns, molecular density, size of the granules, and crowding at the surface. But is this detailed approach strictly required for studying glycogen’s macroscopic properties? In the following, we test various radius values for the spheric description of the glucose units and look whether they affect the macroscopic properties of glycogen.

Three scenarios are investigated: *ρ* = 0 nm, *ρ* = 0.325 nm, and *ρ* = 0.650 nm. For *ρ* = 0 nm, the glucose units have no volume and thus no steric hindrance, allowing chains to overlap. Instead, *ρ* = 0.325 nm is equal to half of the helix’s van der Waals radius, while *ρ* = 0.650 nm is its total radius. By comparing these scenarios, we can evaluate the impact of steric hindrance on the substrate availability during the synthesis of glycogen.

Fig 11 shows that short chains are more abundant for *ρ* = 0 nm than for the other two values of *ρ*. Opposite, long chains are more abundant for non-zero *ρ*. As soon as *ρ* > 0 nm, since elongation involves adding a single glucose unit, while branching means transferring an entire piece of a branch, it is easier to find the necessary space around the substrate for allowing elongation to take place. This is reinforced by the fact that when adding a new glucose unit at the non-reducing end of a branch, we do not elongate the substrate by the total length of a glucose unit, but only its radial contribution, which is *l* = 0.24 nm (see Fig 2). As a consequence, steric hindrance stronger impacts branching than elongation. In other words, the number of branching attempts rejected due to steric hindrance is higher than that of elongation. If we would define effective branching and elongation rates that respectively account for these rejections, the branching effective rate would reduce much more than that of elongation. This would lead to an effective elongation to branching ratio Γ_eff_ which increases with the effect of steric hindrance. In the section Elongation to branching ratio, we concluded that as Γ increases, the overall CLD shifts towards higher DPs and the distribution peak decreases. 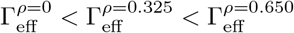 therefore explains both the over-representation of high DPs and the peak reduction as *ρ* increases in Fig 11. Despite these changes in the CLD, as the steric hindrance increases, we can remark that not only is the number of peak conserved (here unimodality) but their location too.

**Fig 11.**
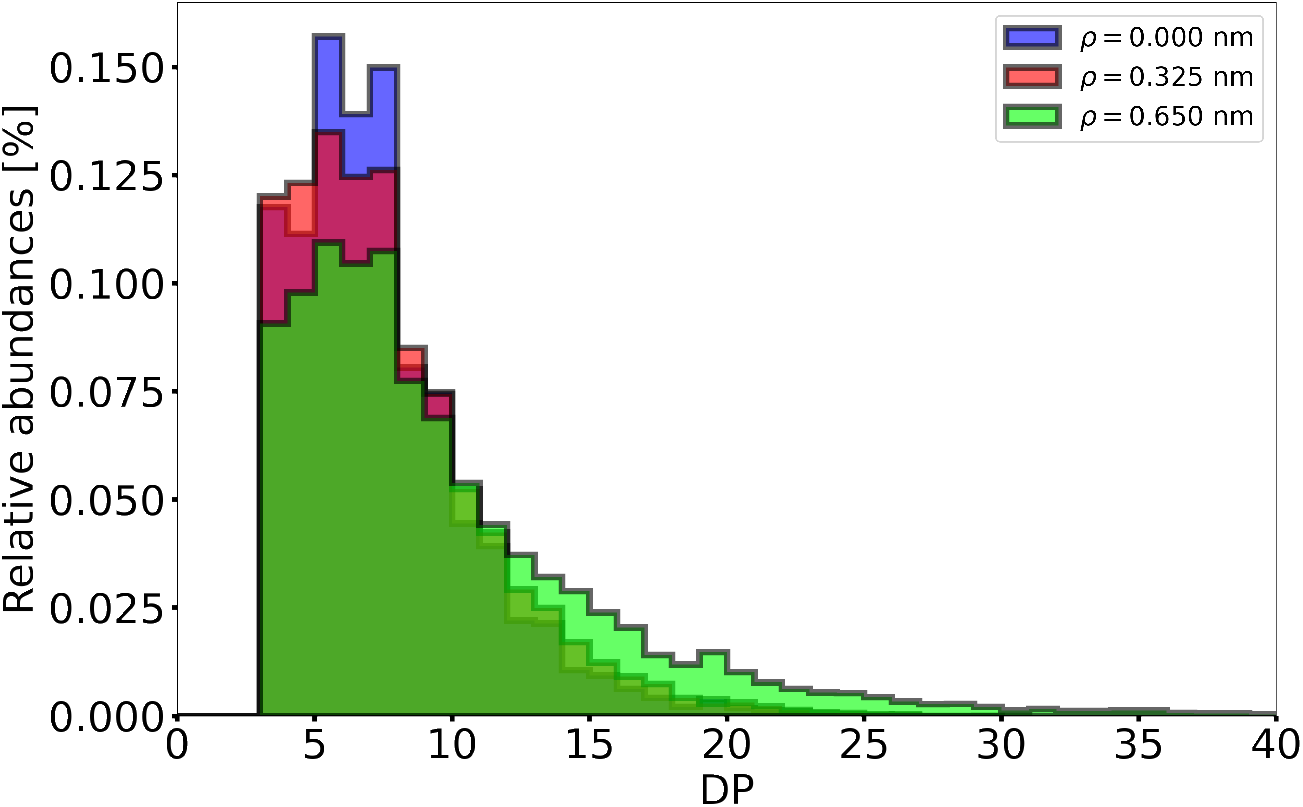
Effect of the steric hindrance on the CLD. Chain length distributions for *ρ* = 0 nm (blue), *ρ* = 0.325 nm (red), and *ρ* = 0.650 nm (green). For *ρ* = 0 nm, the distribution exhibits a higher peak at DP [6–8] than for *ρ* = 0.325 nm and *ρ* = 0.650 nm. Opposite, for *ρ* = 0.325 nm and *ρ* = 0.650 nm, the higher DPs (from DP 12) are over-represented as compare to *ρ* = 0 nm.

### 5.4 Flexible location *versus* strict location branching models

As briefly introduced in the section The model, paragraph Enzymatic reactions, two assumptions for GBE can be made. In the flexible location branching model (used throughout the article) all glucose units that are in the acceptable range can equiprobably be cleaved, and similarly for those that can potentially be transferred. Opposite, in the strict location branching model (introduced here for comparison) GBE always branches at a precise location, i.e. at a given distance from the non-reducing end. For the sake of clarity, these two models and their associated mechanisms are fully detailed in Fig 12. Their patterning impact on the CLD is presented in Fig 13.

**Fig 12.**
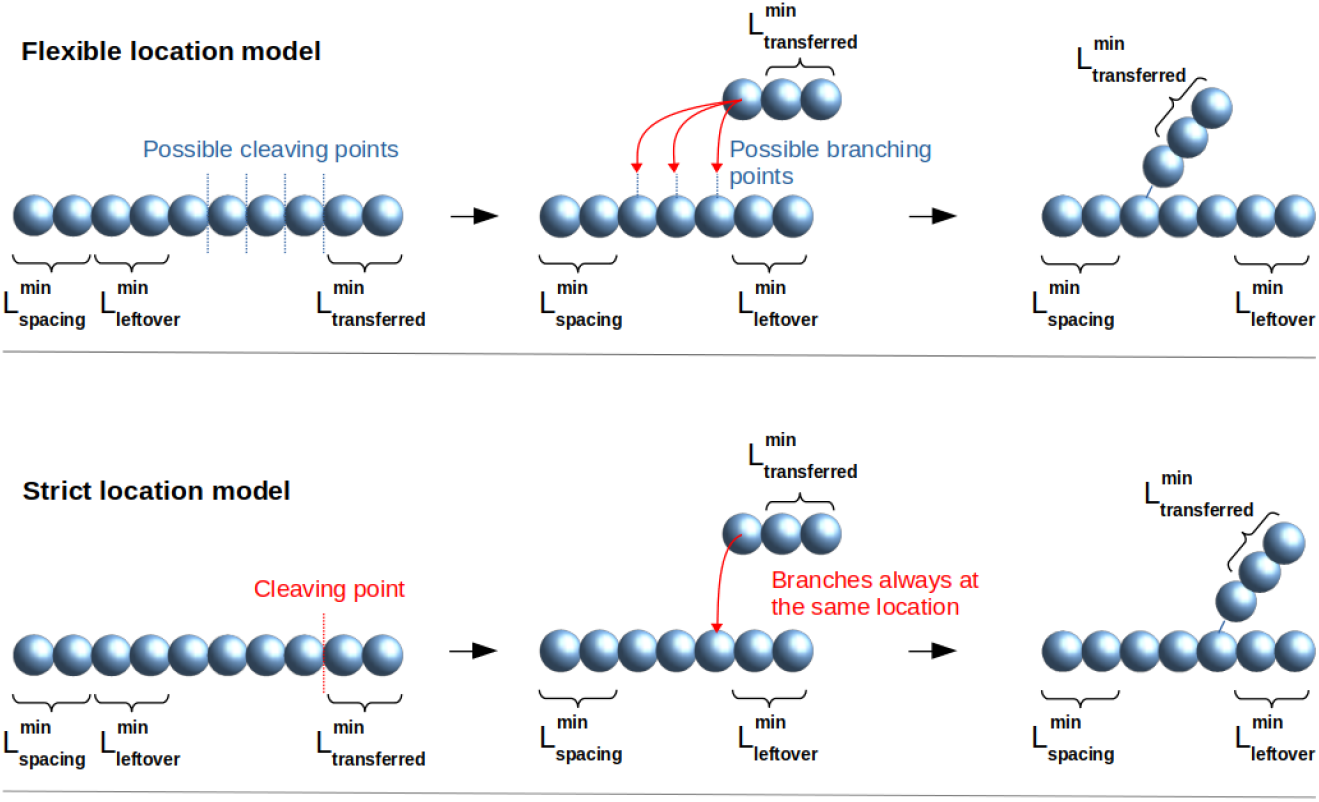
Mechanisms of the flexible location and the strict location branching models. Both models fulfill the minimal lengths requirement (Eq 1). In the flexible location branching model, the cleaving position is randomly picked from a uniform distribution, and so is that of branching. Instead, in the strict location branching model, both the cleaving and the branching always occur at a specific distance from the non-reducing end.

**Fig 13.**
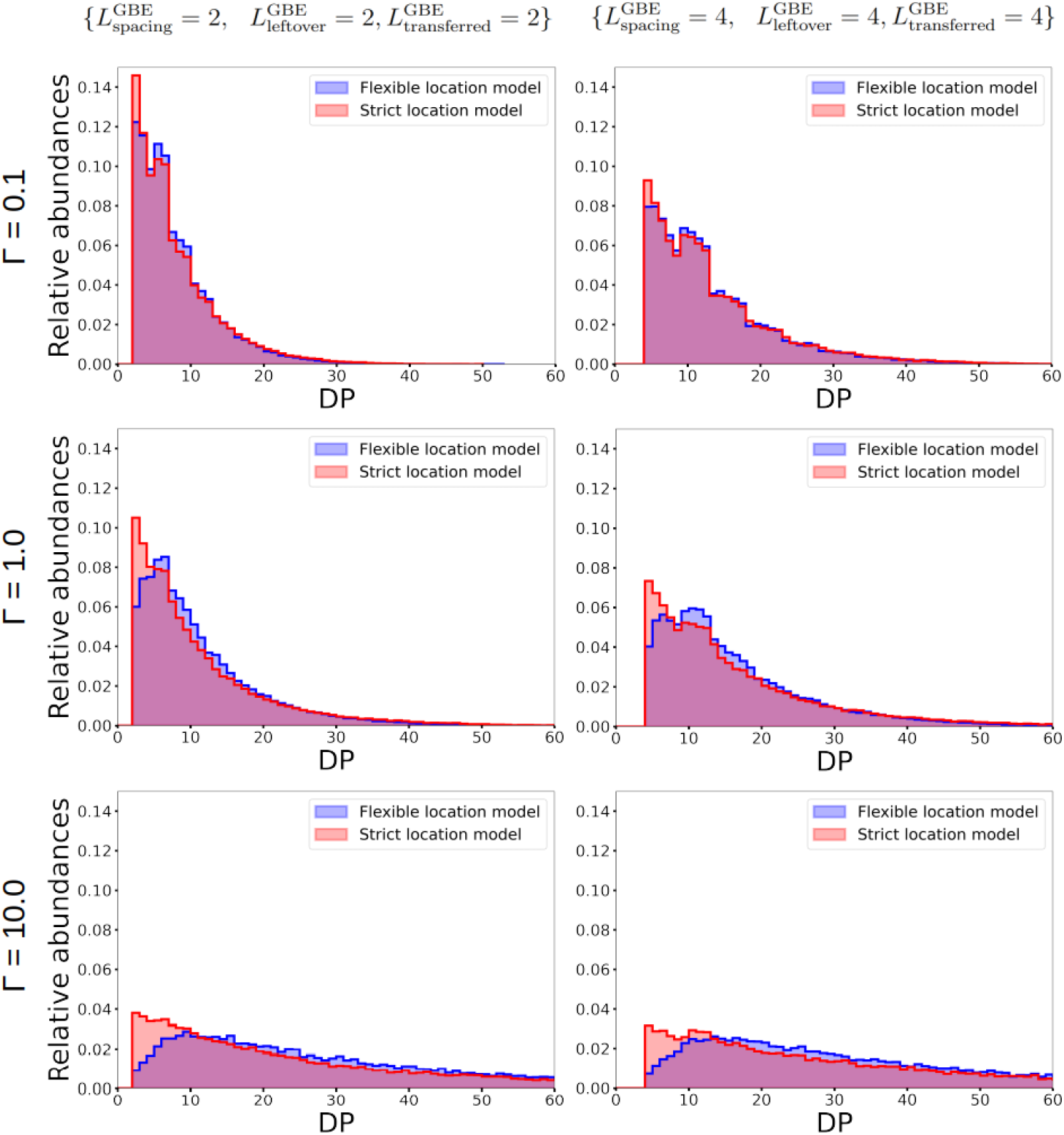
Comparing the impact of the flexible location and the strict location branching models on the CLD. The CLDs for the flexible location and the strict location branching models are shown in blue and red, respectively. Top-left: CLD obtained for small Γ (branching regime), with small minimal lengths for GBE. Top-right: CLD obtained for small Γ (branching regime), with longer minimal lengths for GBE. Increasing the lengths reinforces the multi-modality. Bottom-left: CLD obtained when increasing Γ, with small minimal lengths for GBE. Both conditions contribute to weaken the multi-modality. Bottom-right: CLD when increasing both Γ and minimal lengths for GBE. The first condition tends to weaken multi-modality while the latter one instead reinforces it. As a consequence, multi-modality is observed for the strict location branching model only.

In Fig 13, we compare the two branching models, and vary Γ and 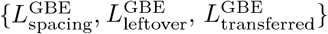, which results in 12 different plots. From the Elongation to branching ratio section, we learnt that increasing Γ spreads the CLD towards higher DPs and reduces the distribution peak typically observed for short chains. In the section Effect of the branching enzyme on the CLD, we observed that increasing the minimal lengths 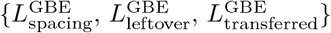 modifies the shape of the CLD that becomes bi- or even multi-modal. When decreasing these lengths, the multi-modality is weakened while it is reinforced by a small Γ.

For Γ = 0.1, the strict location branching model (red) matches the flexible one (blue). That is not surprising, since in this particular regime, when branching strongly dominates over elongation, branching occurs as soon as possible 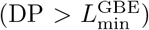, making the two models equivalent.

When Γ increases (second and third lines), the two models are not anymore equivalent. In the strict location model, the length of the transferred branch is always equal to 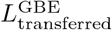, while in the flexible location model, longer chains can be transferred. In both cases, the smallest transferable DP is equal to 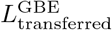 (see section 2.2.3). For the strict location model the latter is the most abundant DP, unlike for the flexible one.

### 5.5 Scope of the parameter space

We apply our fitting procedure to all possible combinations of Γ ∈ {0.1, 0.2, 0.3, 0.4, 0.6, 0.8, 1.0, 2.0, 5.0}, 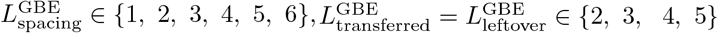 and *ρ* ∈ {0 nm, 0.325 nm, 0.650 nm} while considering two different branching scenarios, i.e. the flexible location and the strict location branching models. It sums up to a total of 1,296 tested parameter sets. The corresponding heat-maps are shown in Fig 14 for the flexible location and Fig 15 for the strict location branching models. In both figures, *ρ* varies across the columns, while 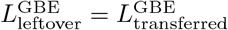 does over the successive rows. On each heatmap, the Y and X axes correspond to 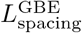 and Γ, respectively.

**Fig 14.**
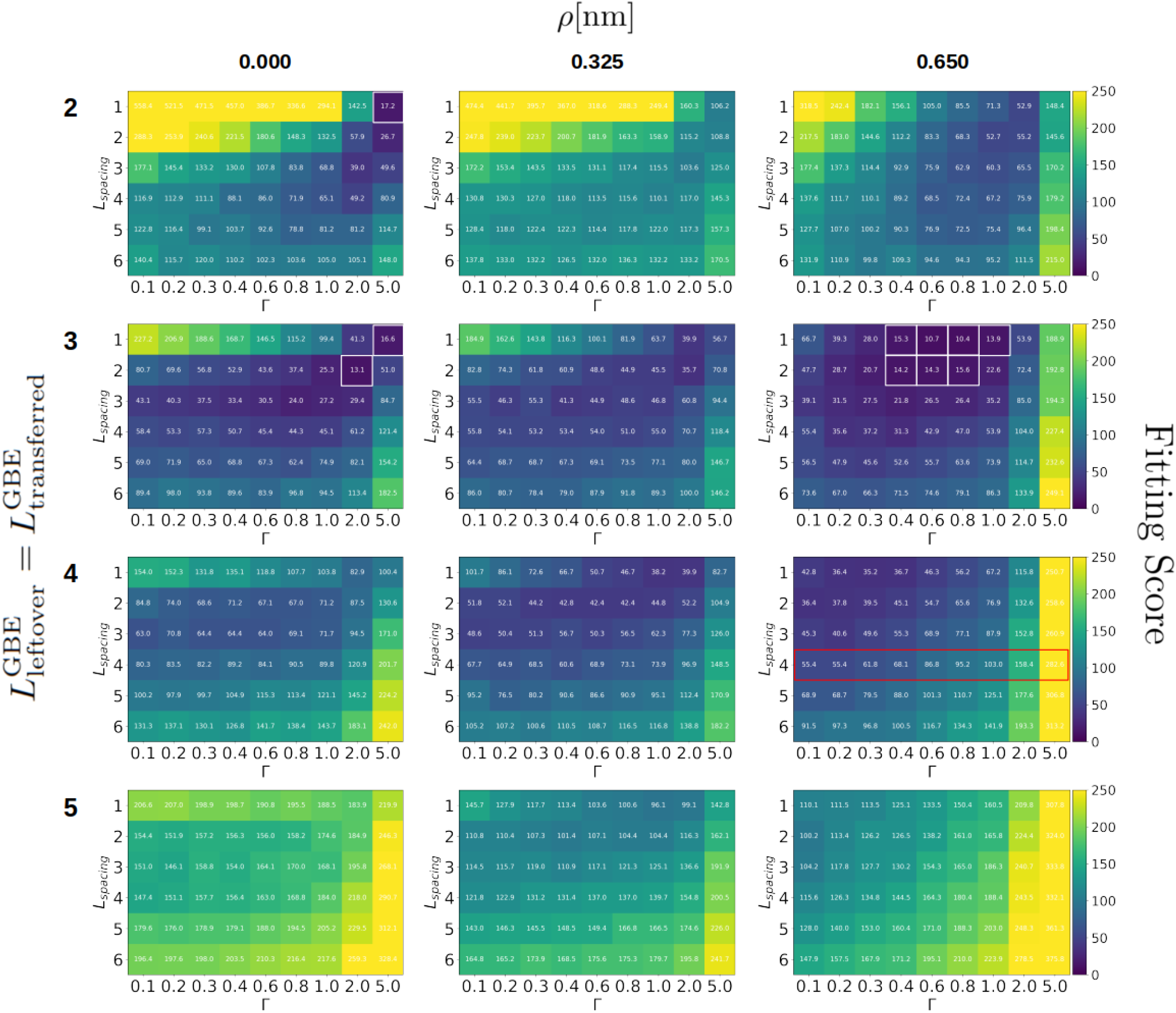
Heat-maps showing fitting scores for the flexible location branching model. The three columns show 3 different radii for the glucose units (*ρ* = 0 nm, 0.325 nm, and 0.650 nm). The 4 rows correspond to distinct values of 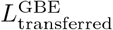 (2, 3, 4, and 5) with 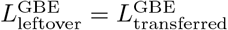. For each heat-map, the values for 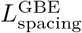 are on the *Y* axis, while the Γ ratio is on the *X* axis. Each cell is characterised by a unique set of parameters 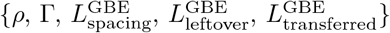. White squares show good scores 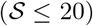, while the red rectangle 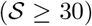 shows the scores obtained for minimal length values as typically assumed in the literature.

**Fig 15.**
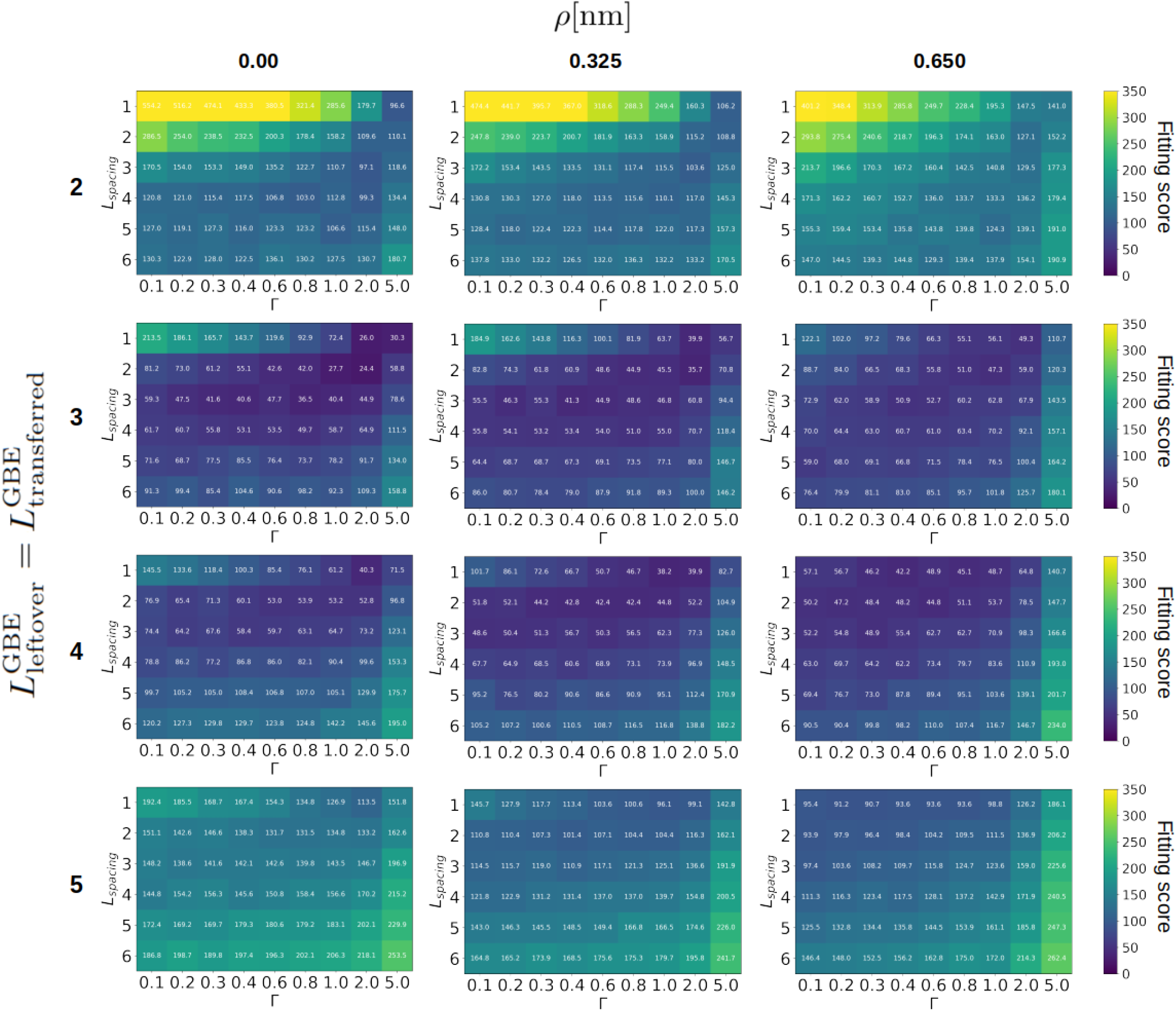
Heat-maps showing fitting scores for the strict location branching model. The three columns show 3 different radii for the glucose units (*ρ* = 0 nm, 0.325 nm, and 0.650 nm). The 4 rows correspond to distinct values of 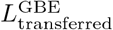 (2, 3, 4, and 5) with 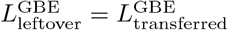. For each heat-map, the values for 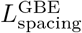 are on the Y axis, while the Γ ratio is on the *X* axis. Each cell is characterised by a unique set of parameters 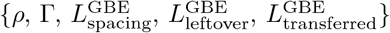. Overall, the scores are much higher than those for the flexible location branching model (see Fig 14), indicating poorer fits. Specifically, no scores below the threshold 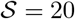 are found.

In Fig 14, all scores below 20 are highlighted with white squares, and correspond to good fits (arbitrary cut-off chosen as up to twice the best-fit). Noticeably, various parameter sets fulfill this criterion and all of them show a small 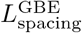 value. As presented in section Comparison to experimental data, paragraph Parameters calibration, the best score is obtained with *ρ* = 0.65 nm (third column). This supports the fact that steric hindrance plays a role in the chain length distribution of real glycogen, although good matches also exist with *ρ* = 0 nm, in which the CLD is not impacted by steric hindrance. When focussing on the good scores (dark blue cells), it appears that changing the elongation to branching ratio Γ can be compensated by varying 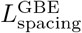, or the two other minimal lengths.

Opposite, when considering parameter values that are typically reported in the literature, we systematically obtain very poor (i.e. high) scores, even upon varying Γ. This case is highlighted in red and discussed in the section Comparison to experimental data, paragraph Parameters calibration. In general terms, for each heatmap, we observe that 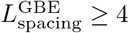 corresponds to poor scores, i.e. 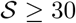. This effect is even more pronounced if 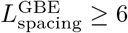, i.e. 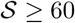 (66.3 and above). Thus, with these parameter sets, our model is not able to reproduce the experimental CLD data by Sullivan and coworkers used for fitting throughout this article [23]. Regarding the enzyme mechanism, this suggests that GBE is able to branch much closer than 4 glucose units away from an existing chain. Similarly, if 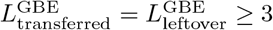, we are not able to fit the experimental data by Sullivan and coworkers. Therefore, these minimal lengths must be shorter than those typically reported in the literature.

In Fig 15, we present analogous results for the strict location model. Overall, the fitting procedure returns much poorer scores than for the flexible location branching model (see Fig 14). This is due to the fact that strict branching locations almost systematically lead to a multi-modal distribution, which is not the case of the experimental CLD data set fitted here [23]. Although multi-modality can be cleared by increasing Γ, this would lead to a reduction of the distribution peak and a shift of the overall distribution towards higher DPs. For the experimental data considered, we could not find a good trade-off, corresponding to good fitting scores. As a result, the strict location model seems to be very unlikely, that is why we instead choose the flexible location branching model for this study.

### 5.6 Probabilistic approach to the A:B ratio

In this section, we describe why the A:B ratio must be equal to 1 when no biological nor biophysical properties of the system are considered, but pure probabilities. As presented in Section 2.3.2, we note as two distinct reactions, branching on either an A or a B chain:

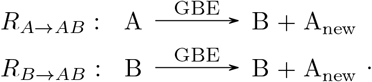

In this purely mathematical framework, A and B chains have the same chance to react, so that the probability for a reaction to occur only depends on the respective number of A (noted *N_A_*) and B (noted *N_B_*) chains. We sketch the associated probability tree in Fig 16. The tree is symmetric, and the number of paths to a given state follows the binomial coefficients. The horizontal dotted line corresponds to *N_A_* – *N_B_* = 0, when there are as many A as B chains. We know that in the example of flipping a coin i times, the central limit theorem tells us that the distribution of the difference in the number of heads and tails, tends to a normal distribution centered in 0 when i tends to infinity. This case corresponds to heads and tails having equal probabilities. In our case, a given state of the probability tree (*N_A_, N_B_*) leads either to the state (*N_A_, N_B_* + 1) with probability 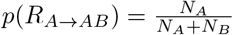, or to (*N_A_* + 1, *N_B_*) with probability 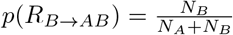. It means that the probability to go from one state to the next one depends on the state of the system, in a way that the probability to come closer to *N_A_ – N_B_* = 0 (horizontal dotted line) is always higher than that of spreading away. Additionally, the distribution of the probabilities remains symmetric with respect to the case *N_A_* = *N_B_*. Based on these considerations, the mean value 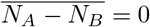 and the distribution is even more peaked than in the simple case of flipping a coin. Since the mean value 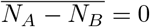, the mean ratio 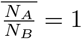, in other words the A:B ratio is 1.

**Fig 16.**
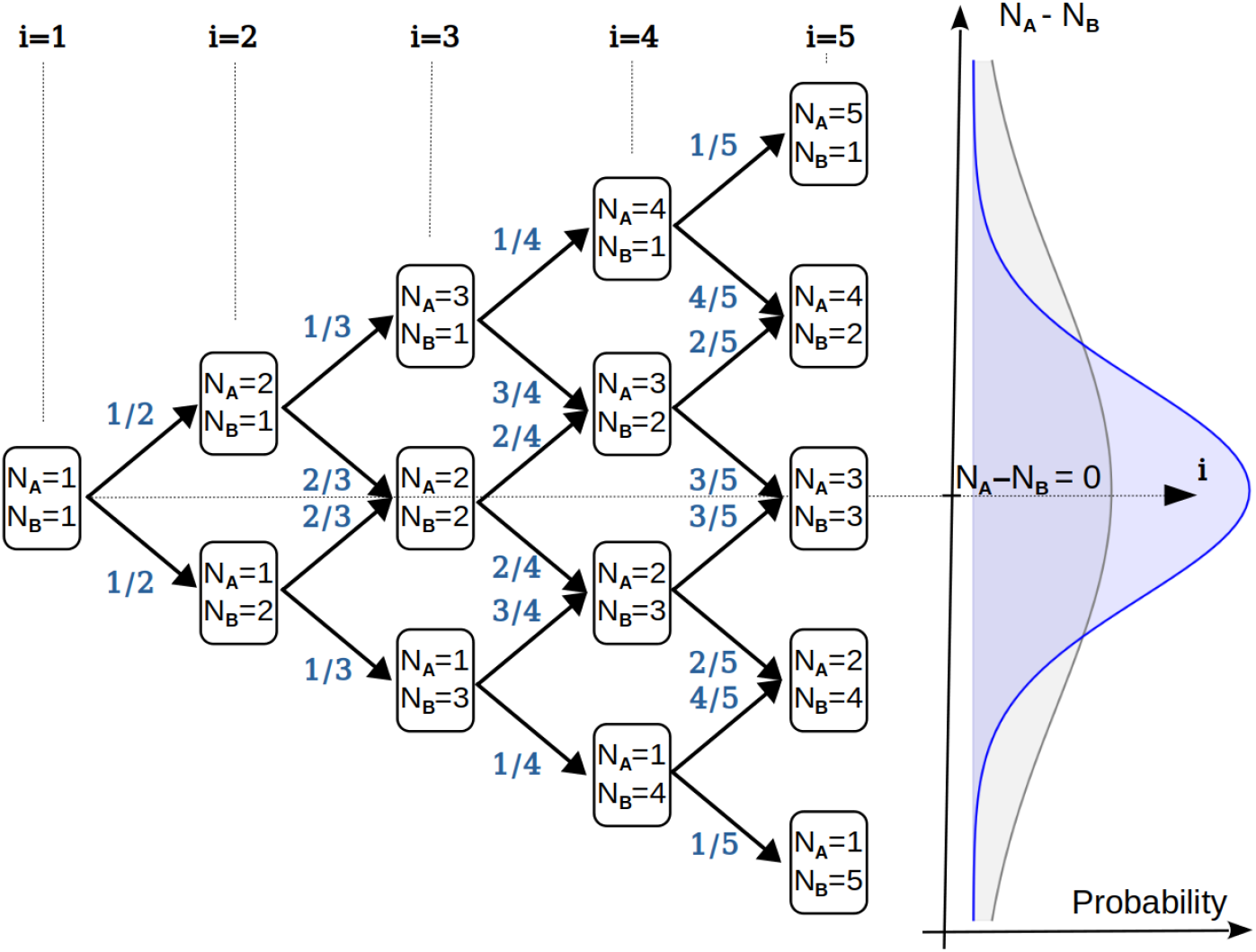
Probability tree of forming a given number of chains of type A (*N_A_*) and B (*N_B_*). Directions that spread the tree are unfavoured, while those oriented towards the center of the tree (reducing the absolute value |*N_A_* – *N_B_*|) are favoured, proportionally to the difference |*N_A_* – *N_B_*|. Therefore, for high numbers of branching reactions (noted i) the distribution (blue) is centered around *N_A_* – *N_B_* = 0, and is thinner than a binomial distribution (grey).

